# *Mycobacterium avium* subspecies *paratuberculosis* targets M cells in enteroid-derived monolayers through interactions with β1 integrins

**DOI:** 10.1101/2024.07.25.604189

**Authors:** Grace Baruta, Kyle L. Flannigan, Laurie Alston, Hong Zhang, Jeroen De Buck, Pina Colarusso, Simon A. Hirota

## Abstract

Paratuberculosis is a global infectious disease caused by the bacterium, *Mycobacterium avium subspecies paratuberculosis* (MAP). MAP infection of ruminants triggers progressive wasting disease characterized by granulomatous lymphadenitis, enteritis, and severe intestinal pathology that often requires early culling of the animal. The resulting economic burden is significant and MAP exposure in the workplace constitutes a significant zoonotic risk. While it has been established the MAP propagates within resident intestinal immune cells, including macrophages and dendritic cells, significantly less is known about how it attaches, enters and traverses the epithelium. The current paradigm suggests MAP infects the small intestinal epithelium by targeting both enterocytes and M cells, with a potential tropism for the latter. In the current study, we employed emerging enteroid technology to identify the target cells for MAP’s entry into the small intestinal epithelium. We generated mouse enteroid-derived monolayers with functional M cells capable of transcytosis. Upon exposure to MAP, the bacteria were detected within both enterocytes and M cells. Following quantification, it was apparent that MAP exhibited tropism for M cells. Complementary studies using the Caco-2/Raji-B co-culture system provided similar results, wherein MAP was found primarily in cells expressing functional M cell markers. Since other mycobacteria have been shown to initiate cell attachment and entry by using a fibronectin-bridging process, we tested whether these interactions were involved in MAP’s targeting of M cells. We found that MAP’s M cell tropism was significantly enhanced in the presence of fibronectin and that this effect was abolished when monolayers were pretreated with an integrin-blocking peptide. Taken together, our data indicate the MAP preferentially targets M cells and that this process involves a fibronectin-bridging process. Furthermore, our data suggest that targeting M cell-associated integrins could provide a mechanism to reduce MAP infection and transmission within livestock herds.

**Author Summary:** In the current study, we sought to determine the target cell for *Mycobacterium avium subspecies paratuberculosis* (MAP), which is the causative agent of Johne’s disease (JD, also termed paratuberculosis) in ruminants. While MAP primarily infects domestic ruminants including cattle, sheep, goats, and deer, it has also been shown to infect wildlife throughout the world, including cats, rabbits, badgers, and wood mice. Given the significant economic burden of MAP infections in livestock, its role in the pathogenesis of JD has been the focus of much research. However, the broad diversity of MAP-susceptible hosts and reservoirs observed calls into question the true scope of MAP infection and transmission and the true number of susceptible hosts. Furthermore, MAP constitutes a zoonotic threat that some have linked to intestinal pathologies, including Crohn’s disease. To date, it is still not known exactly how MAP attaches, enters and traverses the small intestinal epithelium to eventually propagate within resident macrophages and dendritic cells to cause eventual disease. To address this question, we developed a model of the small intestinal epithelium, from mouse enteroids, that contained functional M cells. We found that MAP selectivity enters M cells and that this involves fibronectin-bridging process that targets M cell-associated β1-integrins.

## Introduction

*Mycobacterium avium* subspecies *paratuberculosis* (MAP) is the causative agent of Johne’s disease (JD) in ruminants, a disease commonly transmitted horizontally via the fecal-oral route or vertically via infected milk [1]. Control of MAP transmission of infection is difficult to manage as MAP is shed in the feces well before signs of JD appear [2]. While MAP can multiply within the host, it does so at a relatively slow rate compared to other intestinal bacterial pathogens [3, 4]. MAP has a doubling time of over 22-26 hours and can take several months to grow in culture, depending on the strain and culture conditions [4–6]. Once infected, the current paradigm suggests that several potential fates exist: infection may be cleared by the host; the host may develop subclinical disease; or MAP may eventually trigger a severe intestinal pathology denoted by granulomas, lymphadenitis, enteritis, small intestinal remodelling and diarrhea [7, 8]. Once outward signs of weight loss and diarrhea are present, the animal must be culled, or it will succumb to the infection.

Beyond the economic impact of JD, there is also concern that MAP infection may negatively impact human health [9]. Given the similarities in outward signs of paratuberculosis and Crohn’s Disease (CD), some have posited that MAP may contribute to the pathogenesis of CD. This dates to the early 1900s when two surgeons observed the similarities of MAP infection and CD on the gut epithelium, noting the granulomatous enteritis consistencies in both groups [10–12]. In the early 2000’s, Naser *et al.,* identified MAP in CD patient blood and breastmilk; MAP was found at significantly higher levels in CD patients compared to controls [13–15]. However, it is important to note that MAP has also been found in individuals without CD [14, 16]. To add to this, individuals with occupations that would have high MAP exposure (e.g., farmers, veterinarians) do not report higher incidences of CD [17–19]. While the impact MAP has on CD pathogenesis remains controversial [10, 16], its role as a potential zoonotic pathogen warrants further exploration into how it invades the intestinal epithelium to cause pathology..

Transmission of MAP in ruminants is primarily via the fecal-oral route from infectious animals shedding to susceptible animals through ingestion of MAP in feed, contaminated water beds (including lakes, streams, and other water sources), or through direct ingestion of feces [20]. Infection can also be transmitted from mother to offspring by ingestion of infected milk. Upon ingestion, MAP enters the gastrointestinal (GI) tract and is thought to invade the ileal intestinal epithelium [21]. A seminal study by Payne and Rankin reported that oral MAP infection in calves resulted in lesions within Peyer’s patches (PPs), as noted three months post-infection. This led researchers to suggest a relationship between MAP entry and PPs [22]. As part of the gut-associated lymphoid tissue (GALT), PPs are most commonly found in the ileum of bovines, but are also found in the distal jejunum [22, 23]. The PPs include follicles and domes with an overlaying follicle associated epithelium (FAE) [23], containing specialized membranous or microfold (M) cells, named due to the microvilli on their apical surface [23, 24].

Studies assessing the target cell for MAP entry into the epithelium have provided conflicting results. In an ileal loop ligation model, calves infected with MAP displayed the bacterium within M cells, as assessed by electron microscopy [25]. However, this study only presented images of M cells without including images of any other cell types, with the presence of a single bacterium within a cell to denote infection. Using similar electron microscopy approaches Sigurðardóttir *et al.,* demonstrated the presence of MAP within M following injection of live MAP into the distal small intestine [26]. However, while images of M cells were presented in this report, no other cell types, such as enterocytes or goblet cells, were described. Furthermore, the authors mention that alternate routes of MAP uptake were identified but presented no data to support this statement. Interestingly, in a subsequent study the same group reported that MAP could be cultured from intestinal tissues isolated from regions with and without PPs, suggesting MAP may enter both M cells and enterocytes [27]. In support of these findings, Bermudez *et al.,* demonstrated that MAP could enter and cross the intestinal epithelium in mice devoid of PPs [28]. Furthermore, this study found no significant difference in MAP presence between PPs and non-PP segments of mouse ileum.

Given the inconsistencies present in the literature, in the current study we sought to determine the target cell for MAP’s attachment and entry into the epithelium using emerging enteroid approaches, which constitute primary intestinal epithelial lines generated from crypt stem cells that can be cultured to produce functional M cells. Using broad-reaching outcome measures (confocal microscopy, culture-based approaches) with sufficient resolution, we found that MAP preferentially targets M cells and does so by targeting β1 integrins expressed on the apical surface of these cells. Furthermore, we found that interfering with β1 integrin binding blocked MAP entry into intestinal epithelium, providing a potential target to exploit in the prevention of its spread within livestock.

## Results

### Treating mouse ileal enteroids with RANKL induces the differentiation of M cells

Mouse enteroids were generated as described previously by harvesting crypts from the mouse ileum and culturing them for varying intervals until time of passage (Figure 1) [29]. As anticipated, at day 3 post-passage, enteroids exhibited significant budding, suggesting the induction of differentiation processes (Figure 1C). As days post-passage increased, enteroids accumulated mucous and cellular debris within the lumen (Figure 1 D-F). Due to this, along with an associated decline in viability, all enteroid cultures used in our study were passaged on a weekly basis.

**Figure 1:**
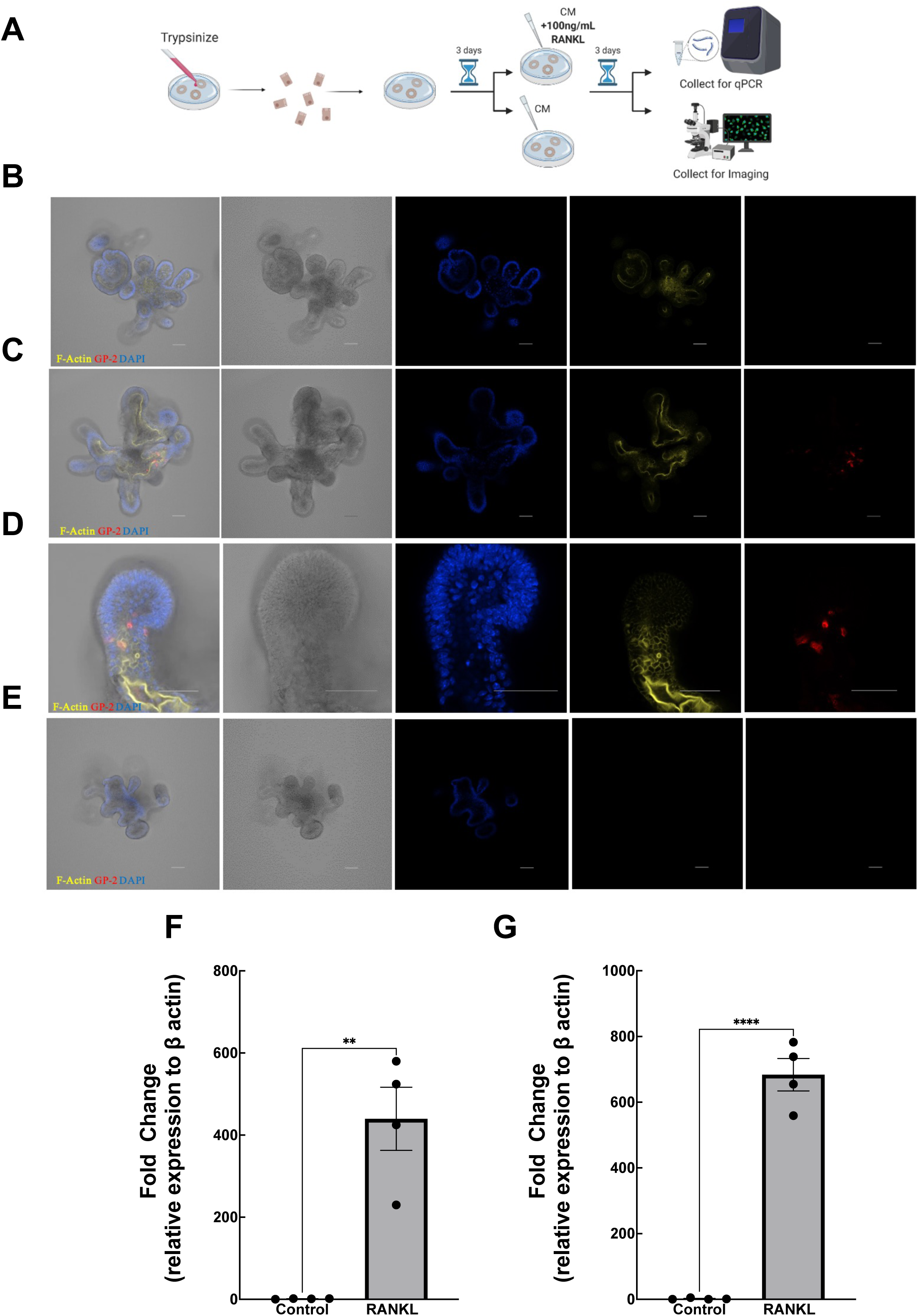
RANKL drives the differentiation of M cells in mouse ileal enteroids. (A-E) Addition of 100 ng/mL RANKL for three days induces GP-2 positive staining in mouse ileal enteroids, as demonstrated by DIC confocal microscopy. Schematic diagram (A) of methods used to generate immunofluorescent images. Ileal enteroids treated with media alone (B) for 6 days lack GP-2 positive signal, while ileal enteroids treated with 100 ng/mL RANKL (C,D) for 3 days display GP-2 positive staining (red). Secondary antibody alone (E) demonstrates the specificity of GP-2 signal in representative immunofluorescent confocal microscopy imaging set. Images and z-stacks were captured on a Nikon-A1R confocal microscope. Blue, 4′,6-diamidino-2-phenylindole (DAPI); yellow, F-Actin; red, GP-2. Data are representative of 5 separate experiments. Scale bar represents 50 µm. (F-G) Ileal enteroids treated with exogenous RANKL express M cell-associated genes. Spi-B (F) and GP-2 (G) expression were significantly increased because of RANKL treatment compared to control. Gene expression was determined by qRT-PCR and is expressed relative to β-actin. Data are expressed as mean ± SEM. * P<0.05, ***P ≤ 0.001 was considered statistically significant as measured via two-tailed Student’s t-test with Prism (GraphPad Software, La Jolla, CA).

Previous reports suggest that the addition of RANKL triggers M cell differentiation in mouse and human enteroid cultures [30, 31]. Given the need to generate enteroids with functional M cells for our studies, we used a previously published protocol, exposing mouse ileal enteroids to RANKL (100ng/mL for 3 days) and assessed differentiation using two markers of M cell differentiation. These included GP-2, which was chosen due to its specificity as a mature M cell marker as well as Spi-B, another standard M cells marker [32]. In our first set of experiments, GP-2 expression was visualized in RANKL- and vehicle-treated ileal enteroids via immunofluorescence staining and confocal microscopy. We also used phalloidin to label organoids because it binds to filamentous actin (F-actin) and thus can be used to examine the morphology of the enteroids. Following 3 days of RANKL exposure, enteroids exhibited increased expression of GP-2 (Figure 1C-D), an effect that was absent in vehicle-treated cultures (Figure 1B). Additionally, RANKL-treated enteroids displayed a significant increase in the expression of *Spi-B* and *GP-2* genes compared to the vehicle treatment group (Figure 1F-G).

These data suggest that exposing 3D enteroid cultures to RANKL increases the expression of M cell differentiation markers.

### Reseeding RANKL-treated 3D enteroids generates 2D monolayers that exhibit the expression of M cell markers

To assess the cellular tropism of MAP in the context of luminal exposure, we sought to generate M cell-containing enteroid-derived monolayers, using a protocol that we reported previously [29]. This approach allows accessibility to both the apical and basolateral compartments for infection experiments to better model the luminal environment. We first seeded enteroid-derived monolayers onto transwells and treated with RANKL for 3 days (100 ng/mL; new media containing RANKL added daily to the basolateral compartment of the transwell). Surprisingly, this approach did not induce the expression of M cell marker GP-2, as assessed by immunofluorescence staining and confocal microscopy (Figure 2A-B). Next, we treated 3D enteroids with RANKL for 3 days, then seeded them onto as monolayers. This approach resulted in monolayers exhibiting GP-2 expression (Figure 2C). We also observed a significant increase in *Spi-B* and *GP-2* gene expression in monolayers generated using this approach (Figure 2E-F). Taken together, the findings suggest that, under our current culture conditions, RANKL-induced M cell differentiation only occurs in 3D enteroids, but that reseeding 3D organoids is a viable approach to generate monolayers expressing M cells markers.

**Figure 2:**
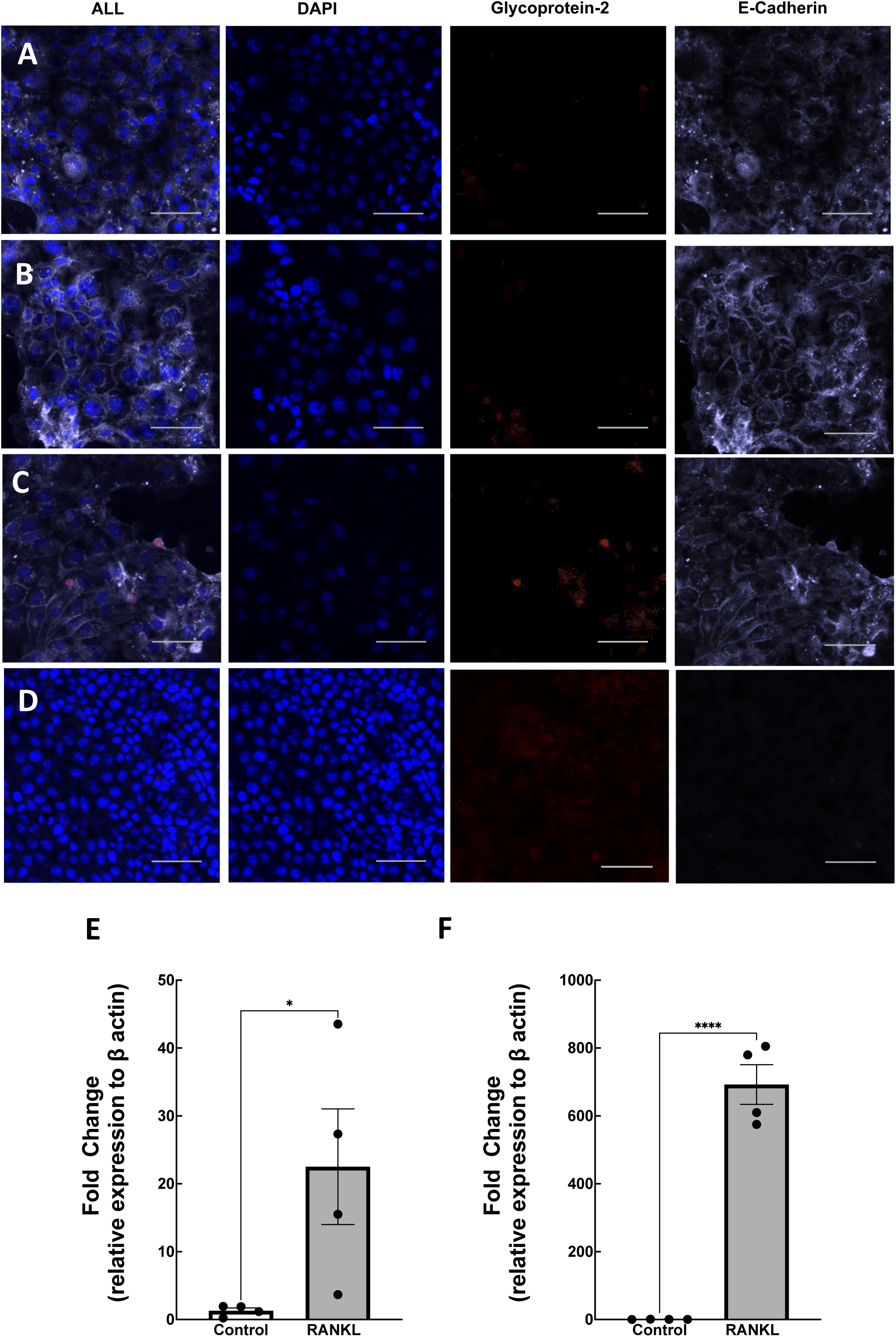
Ileal enteroid-derived monolayers treated with exogenous RANKL prior to seeding results in the differentiation of M cells as indicated by GP-2 positive staining and Spi-B and GP-2 gene expression. Mouse enteroid-derived monolayers were treated with media alone (A) or 100 ng/mL RANKL post-seeding (B). Monolayers treated with media alone or RANKL post-seeding did not display any GP-2 positive staining. However, RANKL-treated 3D enteroids and the monolayers generated from them (C) display GP-2 positive staining (red). Secondary control (D) shows specificity of staining. Representative confocal images from 5 unique experiment sets. Images and z-stacks were captured on a Nikon-A1R confocal microscope. Blue, DAPI; purple, ECAD; red, GP-2. Scale bar represents 50 µm. To evaluate gene expression, RANKL-treated and control monolayers were collected, and Spi-B (E) and GP-2 (F) gene expression evaluated. Gene expression was determined by qRT-PCR and is expressed relative to β-actin. Data are shown as means ± SEM from 4 experiments. * P<0.05, ****P ≤ 0.0001, was considered statistically significant as measured via two-tailed Student’s t-test with Prism, n=4 (GraphPad Software, La Jolla, CA).

### RANKL treatment induces the differentiation of active M cells in mouse 2D enteroid-derived monolayers

The presence of GP-2 is restricted to mature M cells capable of transcytosis, suggesting that the M cells in our system are functional [33]. We next sought to verify that the M cells in our monolayers exhibited functional transcytosis. As a direct measure of functional M cells, inert bead uptake was used to measure active transport, as reported previously [31]. In brief, 1×10^5^ 1µm fluorescent polystyrene beads were added to the apical side of 2D enteroid-derived monolayers and incubated for 1 hour and collected for immunofluorescent confocal imaging. We found that the fluorescent polystyrene beads co-localized with cells expressing GP-2, the mature M cell marker (Figure 3A-B). Confocal z-stack assessment demonstrated that these beads were within the M cells of the monolayer (1μm beads [red] and GP-2 positive M cells [green]; Figure 3B). To evaluate these data quantitatively, the percentage of enterocytes (E-cadherin positive [ECAD]) or M cells exhibiting positive bead uptake were enumerated. The percentage of M cell-associated beads was significantly higher compared to enterocyte-associated beads (Figure 3C). Additionally, our findings reveal a significant increase in beads found in M cells compared to enterocytes, normalized to cell count (Figure 3D). Together, these data suggest that the M cells induced by RANKL treatment exhibit the appropriate functionality (i.e., active particle uptake) for use in our subsequent host-pathogen interrogation experiments.

**Figure 3:**
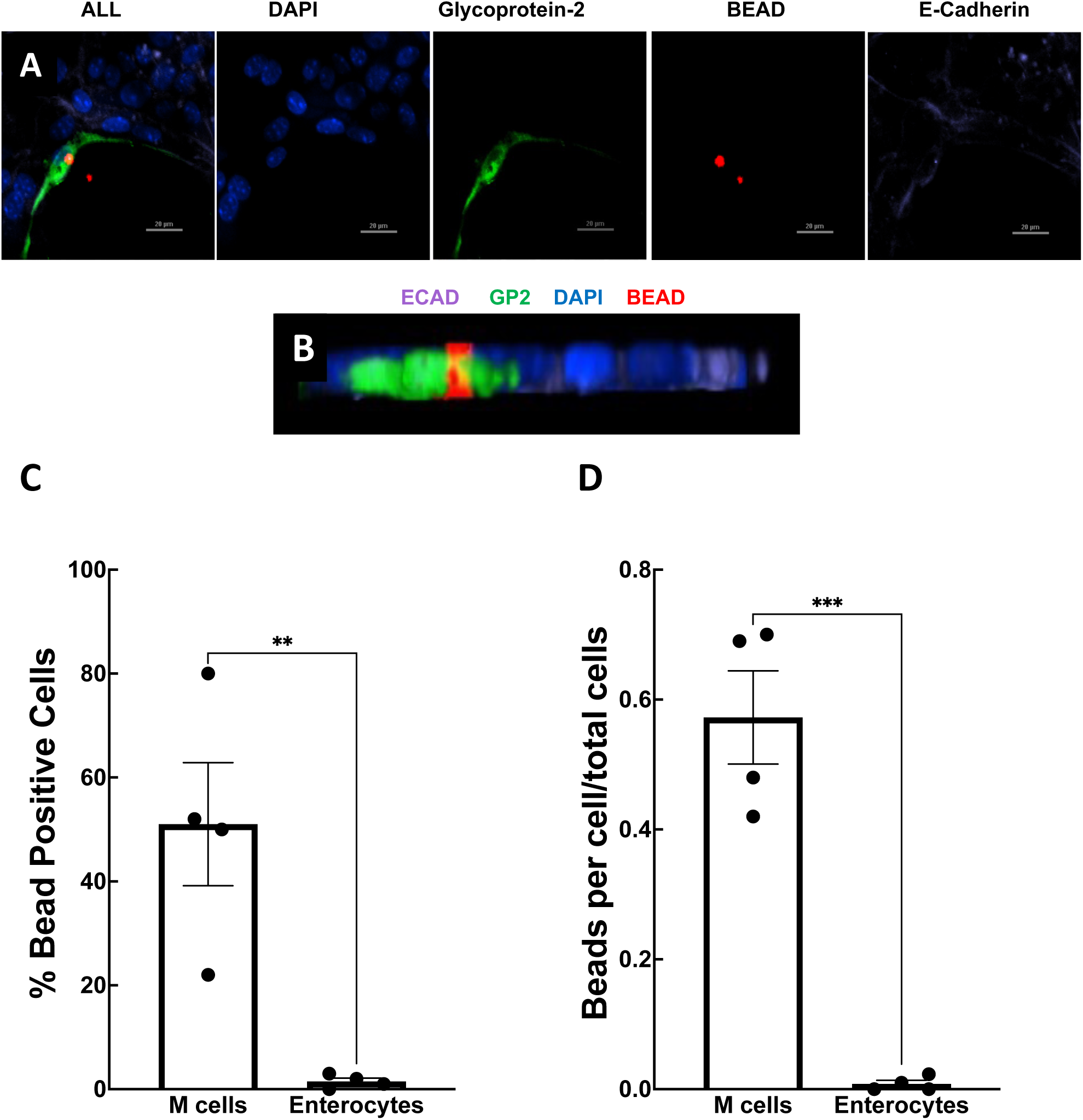
RANKL-treated enteroid-derived monolayers demonstrate functional M cells through uptake of inert fluorescent microparticles. RANKL-treated, ileal enteroid-derived monolayers were exposed to 1 µm fluorescent-Fluorospheres for one hour and collected for immunofluorescence confocal imaging. Representative image demonstrates bead and M cell positive staining co-localization (A) with GP-2 (green), fluorescent fluorosphere (red), ECAD (purple) and DAPI (blue). Confocal microscopy z-stack 3D rendering of representative image in (A) demonstrates colocalization of fluorescent 1 μm microparticle and GP-2 positive stained M cells (B). Images and z-stacks were captured on a Nikon-A1R confocal microscope and are representative of 4 experiments. Scale bar represents 20 µm. Percent epithelial cell positive beads (C) and cell-associated beads (D) were calculated from three images per well, three wells per group, and 4 separate experiments. Data are shown as means ± SEM. ** P≤ 0.01, ***P ≤ 0.001 was considered statistically significant as measured via two-tailed Student’s t-test with Prism (GraphPad Software, La Jolla, CA).

### MAP preferentially targets M cells in enteroid-derived monolayers

Using our model of enteroid-derived monolayers containing functional M cells, we next sought to assess the cell tropism of MAP. To understand the kinetics of cellular invasion, 5×10^5^ CFU/mL GFP-MAP was added to monolayers and incubated for 30 minutes, 1 hour, or 6 hours; these timepoints were based on previous studies performed in cell line-based experiments [34]. We found that MAP translocated across the monolayer quickly, as we observed GFP-MAP signal present in monolayers at 30 minutes (Figure 4A) post-exposure, but this signal was absent at later timepoints (Figure 4B-C). Therefore, moving forward we chose the 30-minute timepoint to evaluate MAP uptake within the intestinal epithelium.

**Figure 4:**
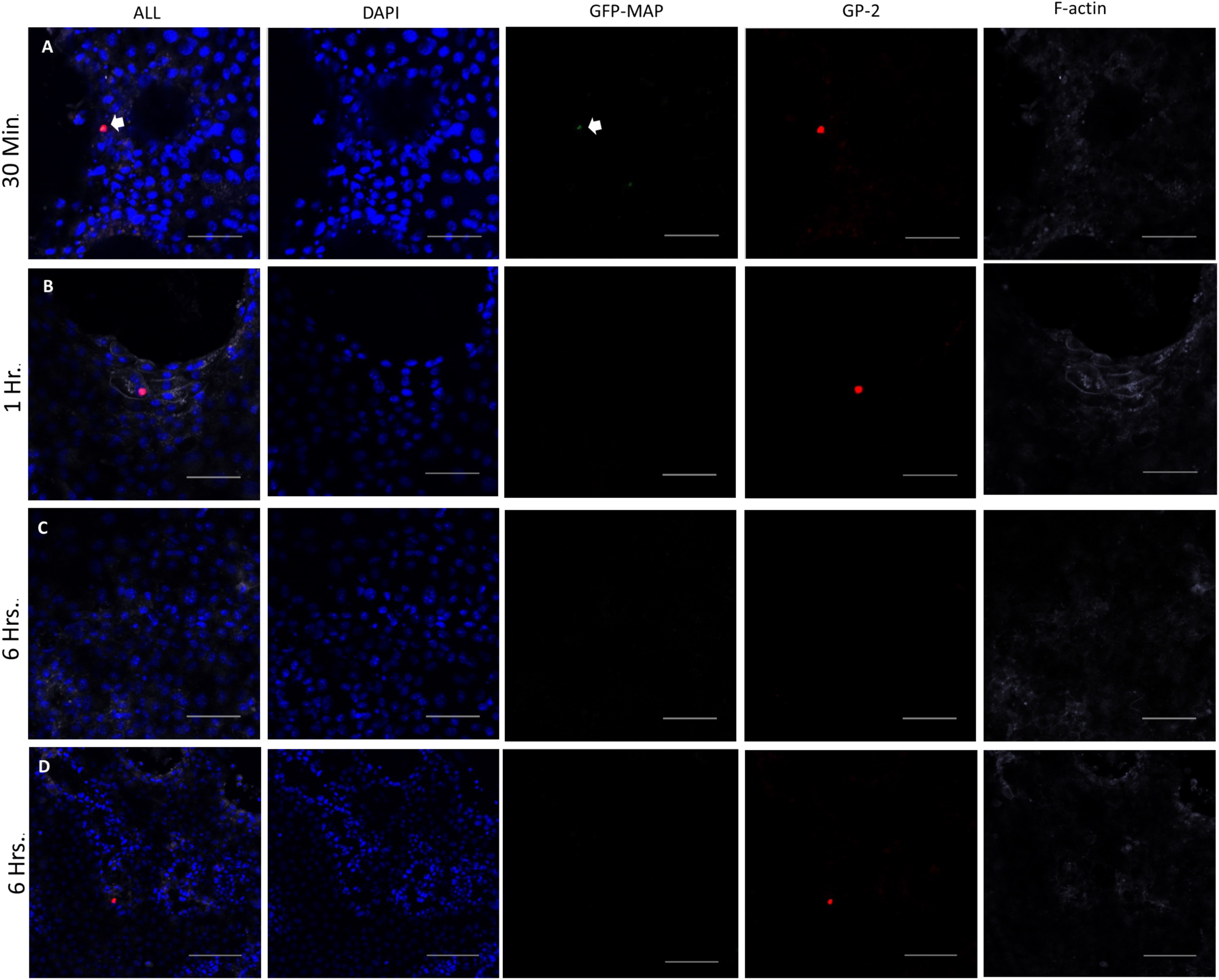
MAP can be seen within the enteroid-derived monolayers at 30 minutes post MAP exposure, but not evident at 1- or 6-hours post-exposure. RANKL-treated, ileal enteroid-derived monolayers were exposed to 5×10^5^ CFU/mL GFP-MAP for 30 minutes (A), 1 hour (B), or 6 hours (C, D), fixed, and stained for GP-2 (red), F-actin (purple), and DAPI (blue). GFP-expressing MAP can be seen at the 30-minute timepoint (A) but is not seen at the 1-hour (B) or 6-hour (C, D) timepoints. Images are representative of 3 separate experiments, with scale bar representing 50 µm (A-C) and 100µm (D). Images and z-stacks were captured on a Nikon-A1R confocal microscope.

Next, to the test the hypothesis that MAP selectively targets M cells for entry, enteroid-derived monolayers were incubated with 5 x 10^5^ CFU/mL GFP-MAP for 30 minutes. Immediately following GFP-MAP exposure, monolayers were fixed and stained for GP-2 (M cells) and ECAD (enterocytes), and the samples were images using confocal microscopy. Images were subsequently quantified in a blinded manner to determine the number GFP-MAP-positive M cells versus enterocytes. Our imaging revealed that GFP-MAP could be found within M cells (Figure 5A-C) and enterocytes (Figure 5D-F). However, quantification revealed that MAP favoured entry into M cells, as indicated by a significant increase in the percentage of GFP-MAP-positive M cells (Figure 5G-H). Furthermore, adherent GFP-MAP (GFP signal not coincident within the monolayer plane as assessed by visual inspection, but associated with apical aspect of the cells) was significantly higher on M cells compared to enterocytes (Figure 5I). Taken together, these data support our hypothesis that M cells are the major cellular target for MAP attachment and cellular invasion.

**Figure 5:**
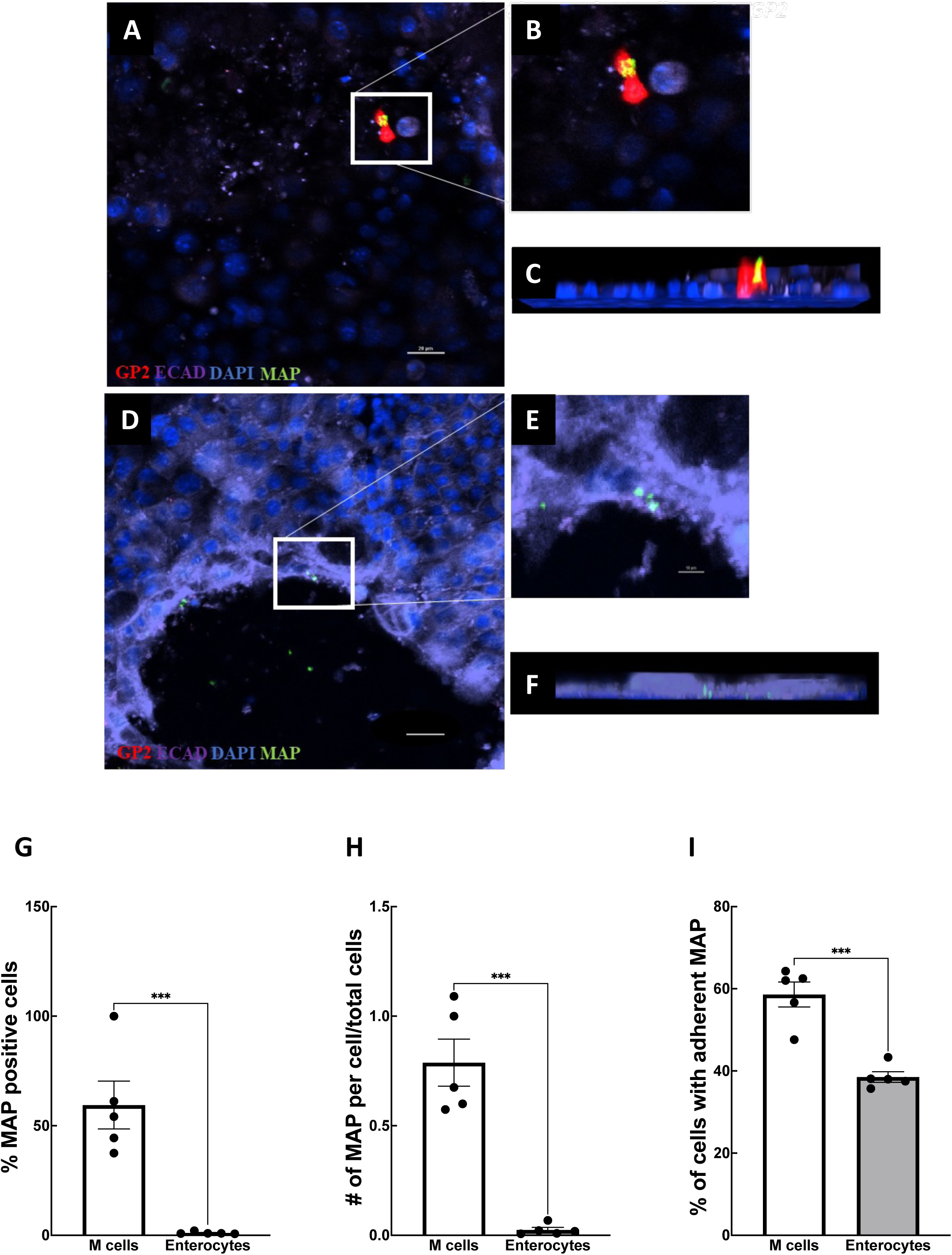
MAP displays tropism for M cells. RANKL-treated ileal enteroid-derived monolayers were exposed to 5×10^5^ CFU/mL GFP-MAP for 30 minutes, fixed, and stained for GP-2 (red), ECAD (purple), and DAPI (blue). GFP-expressing MAP colocalizes with M cell marker, GP-2, (A) as demonstrated at an increased magnification in (B) and in z-stack rendering (C). GFP-expressing MAP colocalizes with enterocyte marker, ECAD (D), as shown at an increased magnification in (E) and z-stack rendering (F). Images are representative of 5 separate experiments, with scale bar representing 20 µm (A, F) and 10µm (E). Images and z-stacks were captured on a Nikon-A1R confocal microscope. Percent MAP positive M cells were significantly higher compared to percent MAP+ enterocytes (G). MAP numbers were significantly increased in M cells compared to enterocytes, normalized to cell type (H). Adherent MAP counts were significantly higher in M cells compared to enterocytes (I). Statistical tests used were two-tailed Student’s t-test, N=5 (***P ≤ 0.001) with values representing mean ± SEM, with the Shapiro-Wilk test performed to assess normal distribution.

### Caco-2/Raji-B co-cultures exhibit functional M cells

To support our findings from the mouse ileal enteroid-derived monolayer experiments, we sought to use a complementary *in vitro* system involving the co-culture of human Caco-2 cancer epithelial cell line with lymphoblast-like Raji-B cells, as previously described [35]. In mono-culture, Caco-2 cells form a homogenous, confluent, and impermeable monolayer, whereas co-culture with Raji-B cells triggers the differentiation of M cell-like cells. We first confirmed the induction of sialyl Lewis A antigen (SLAA; a human M cell marker [36]) expression in the context of Caco-2/Raji-B co-culture (Figure 6). Next, we evaluated the ability of SLAA-expressing M cell-like cells to take up microparticles. To do so, we performed fluorescence imaging on the mono- and co-culture systems that were incubated with 1 µm fluorospheres for 1 hour [31]. We then fixed the cells and imaged the samples using confocal microscopy. We quantified bead translocation via measuring the fluorescent signal within the basolateral chamber compared to a standard curve. Using confocal microscopy, we were able to observe the presence of beads within the SLAA-positive staining regions (Figure 7A-B) of our co-culture model system (Figure 7C). The z-stack rendering in the representative image below shows a bead directly co-localized with the SLAA-positive cells (Figure 7D); an adjacent bead is found to not be within the cell boundary (Figure 7E). Quantification of translocation revealed that SLAA-expressing Caco-2/Raji-B co-cultures exhibited significantly more bead accumulation within the basolateral compartment when compared to the Caco-2 mono-cultures (Figure 7F); while both systems exhibited equivalent measures of barrier function (TEER measurements; Figure 7G). Taken together, these data suggest that the Caco-2/Raji-B co-culture system contains functional M cell-like cells that exhibit the ability to facilitate transcytosis and can be used in subsequent MAP invasion studies.

**Figure 6:**
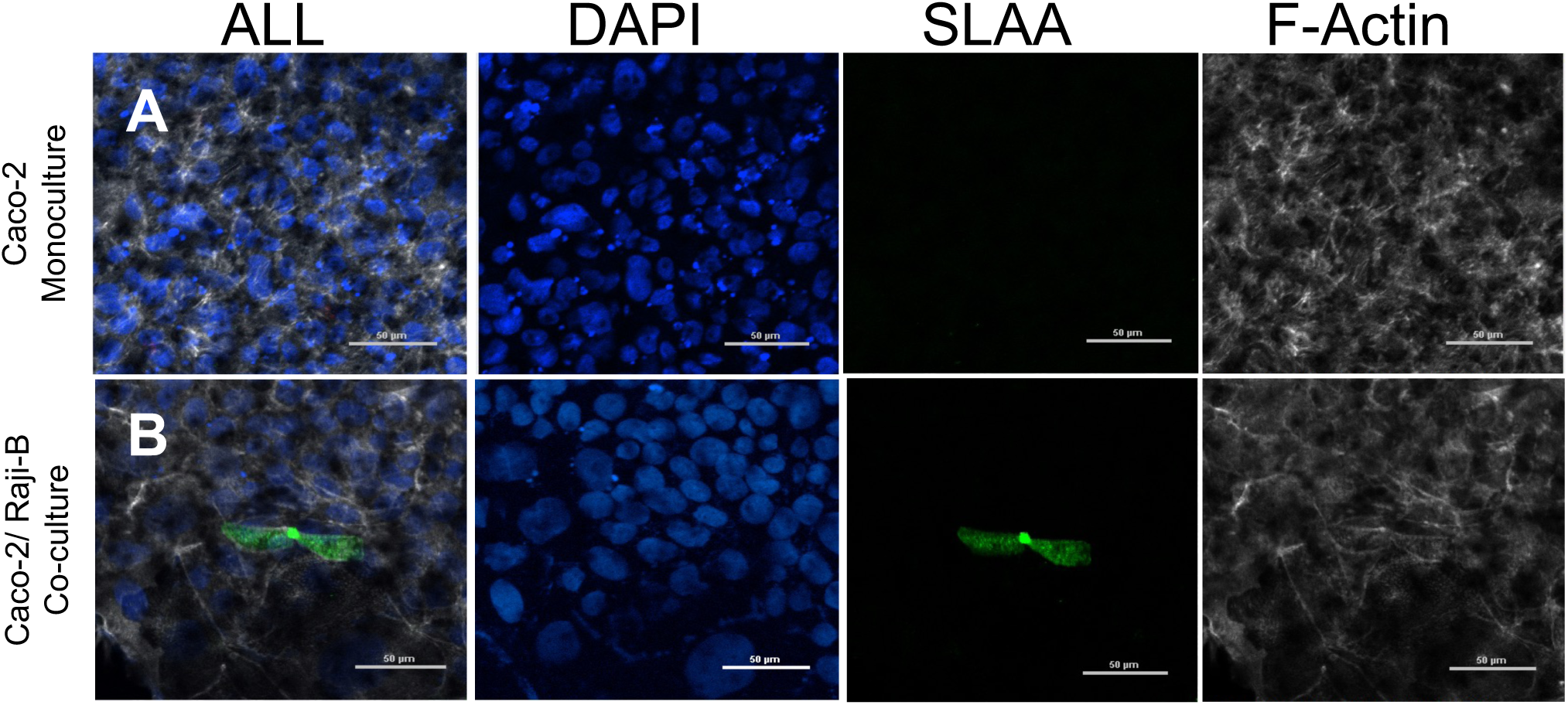
Establishment of the Caco-2/Raji-B co-culture model system. SLAA positive staining (green) was not observed in monoculture (A), but was present in the co-culture system (B). Images and z-stacks were captured on a Nikon-A1R confocal microscope. Scale bar represents 50 µm. DAPI (blue), SLAA (green), F-actin (white). Representative confocal images from 3 separate experiments.

**Figure 7:**
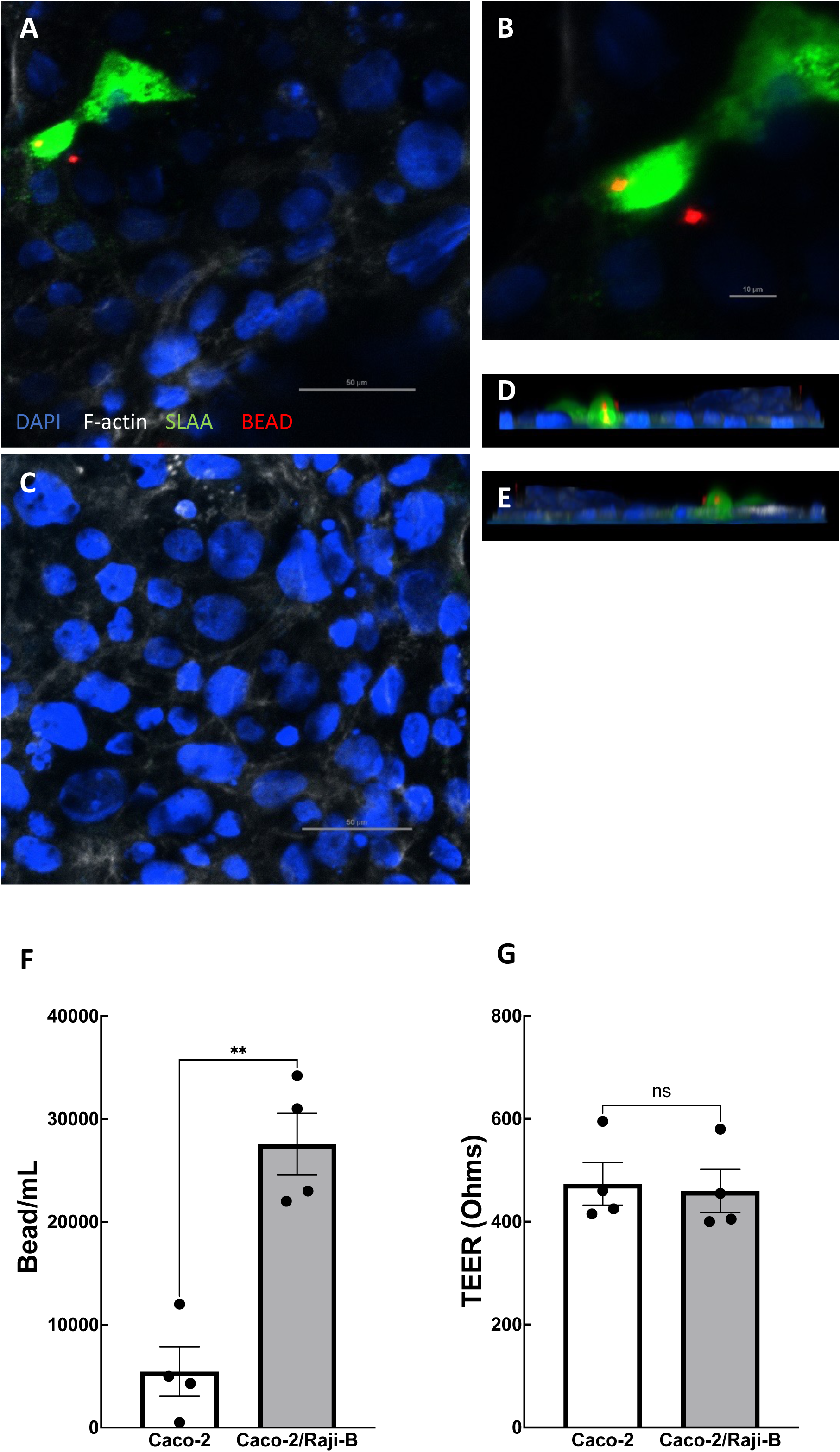
Transport of fluorospheres across Caco-2 monocultures and Caco-2/Raji-B co-cultures. Mono- and co-cultures were incubated with fluorospheres for 1 hr at 37°C and monolayers collected for immunofluorescence imaging to evaluate bead uptake. Bead uptake (red; A) was observed within M cell-like cells (SLAA stain; green) as seen in the magnified image (B) and z-stack cross-section (D, E) of the co-culture system. Representative confocal image of mono-culture (C) did not have beads evident within the monolayer. To evaluate the bead adjacent to SLAA stain, the z-stack cross-section (E) demonstrates that the bead is not co-localized with either F-actin (white) or SLAA (green). Images and z-stacks were captured on a Nikon-A1R confocal microscope. Scale bar represents 50µm (A, C) and 10µm (D). Both culture systems exhibited a similar degree of barrier function, as assessed by TEER measurements (corrected to baseline TEER of Matrigel-coated Transwell; F). To evaluate translocation of beads, SpectraMax i3x Multi-Mode Microplate Reader was used in fluorescence intensity mode to evaluate translocated beads compared to a standard curve to measure beads per mL Representative images from 4 separate experiments, 4-8 wells per group (G). Data are shown as means ± SEM. ** P<0.01 was considered statistically significant as measured via two-tailed Student’s t-test with Prism (GraphPad Software, La Jolla, CA).

### SLAA-positive M cell-like cells are preferentially targeted by MAP

Since our Caco-2/Raji-B co-culture system exhibited functional M cell-like cells, we next tested whether MAP would preferentially invade these SLAA-positive cells. To do this, GFP-MAP was added to both Caco-2 mono-culture and Caco2/Raji-B co-culture systems for 1 hour and monolayers were harvested for several downstream assays. To evaluate epithelial cell targets of MAP, monolayers were collected for immunofluorescence imaging. Representative images and z-stacks are displayed in Figure 8; an apical view with SLAA-positive staining can be seen in Figure 8A. As SLAA staining is restricted to the apical surface of this co-culture system, the bottom of the z-stack close to the membrane is displayed in Figure 8B, where GFP-MAP can be seen directly under the SLAA staining present in Figure 8A. To visualize this together, z-stacks are displayed in Figure 8C-E, where green GFP-MAP signal is evident directly under red SLAA staining (Figure 8D-E). GFP-MAP was also evident in areas devoid of apical SLAA-positive staining (Figure 8F). This demonstrates that GFP-MAP is found within both M cell-like cells and enterocytes of the Caco-2/Raji-B co-culture system.

**Figure 8:**
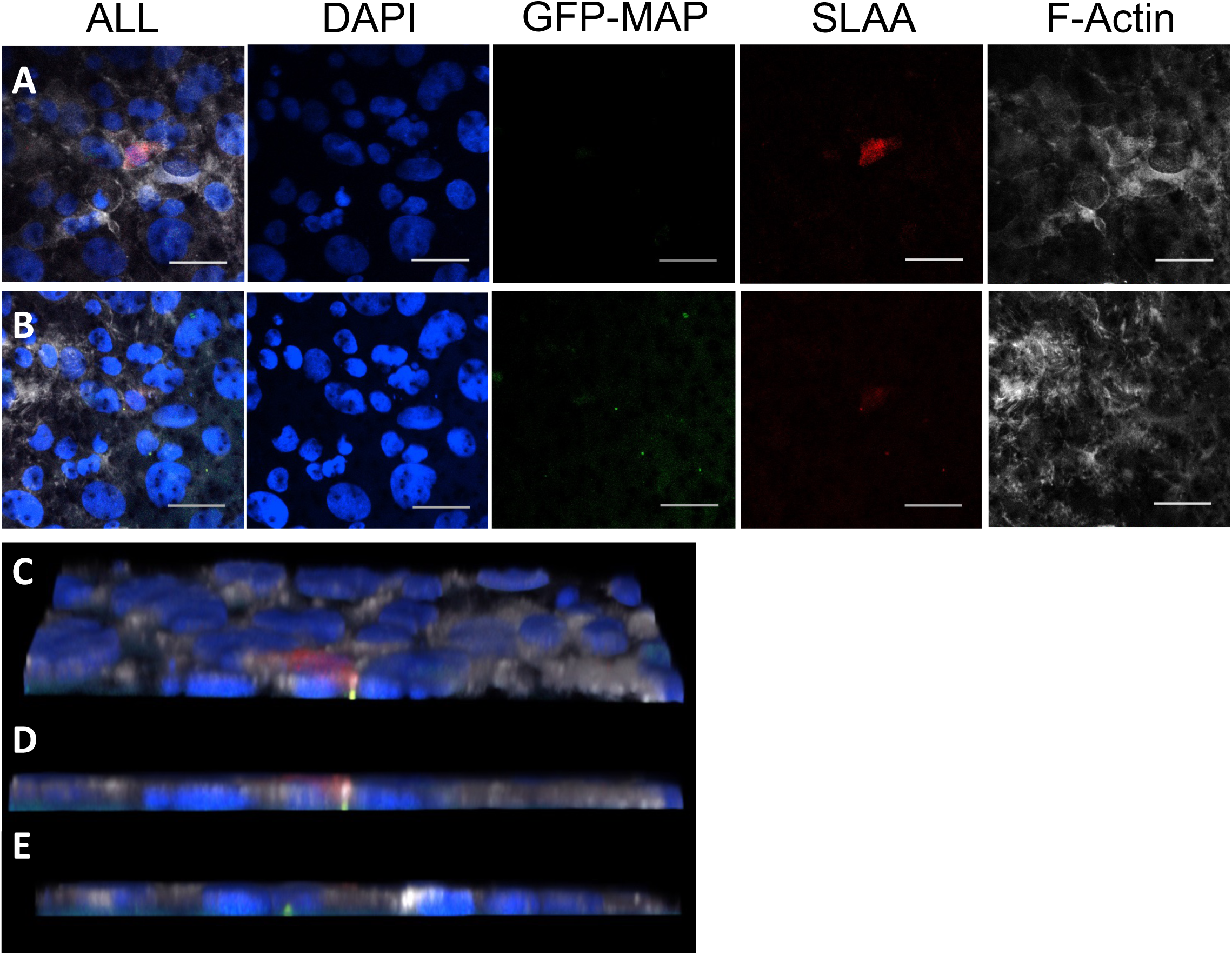
GFP-MAP is found within M cell-like cells and enterocytes in co-culture model system. The Raji-B/Caco-2 co-culture apical view displays the presence of M cell marker SLAA (red) (A), which is apically restricted and observed at the top of the representative z-stack (C, D). Closer to the Transwell membrane (B, D), GFP-MAP is visible directly below apical SLAA staining, indicating the presence of MAP within this SLAA-positive M cell-like cell. GFP-MAP can also be seen in areas not covered with SLAA staining (E), indicating the presence of MAP uptake in enterocytes. Representative images from 4 separate experiments, scale bar represents 50 µm. Images and z-stacks were captured on a Nikon-A1R confocal microscope.

To evaluate whether the presence of SLAA-positive M cell-like cells enhanced MAP uptake, we quantified its intracellular presence using culturing approaches as described by others previously [37–39]. GFP-MAP was added to the apical chamber of the monolayer culture systems for 1 hour. After this, the basolateral chambers were collected for fluorescence measurements and the monolayers washed with PBS, subsequently treated with gentamycin, cells lysed, and serial dilutions plated for CFU quantification (workflow depicted in Figure 9A). Both CFU counts and the quantification of GFP-MAP fluorescence showed that Caco-2/Raji-B co-cultures exhibited significantly more internalized MAP than Caco-2 mono-cultures (Figure 9B-C). The enhanced MAP internalization and translocation across the Caco-2/Raji-B monolayers occurred in the absence of any differences in TEER, indicating comparable barrier integrity in both culture systems (Figure 9D). Taken together, these data align with those generated in our enteroid-derived monolayers, wherein M cells are the predominant target of invasion for MAP.

**Figure 9:**
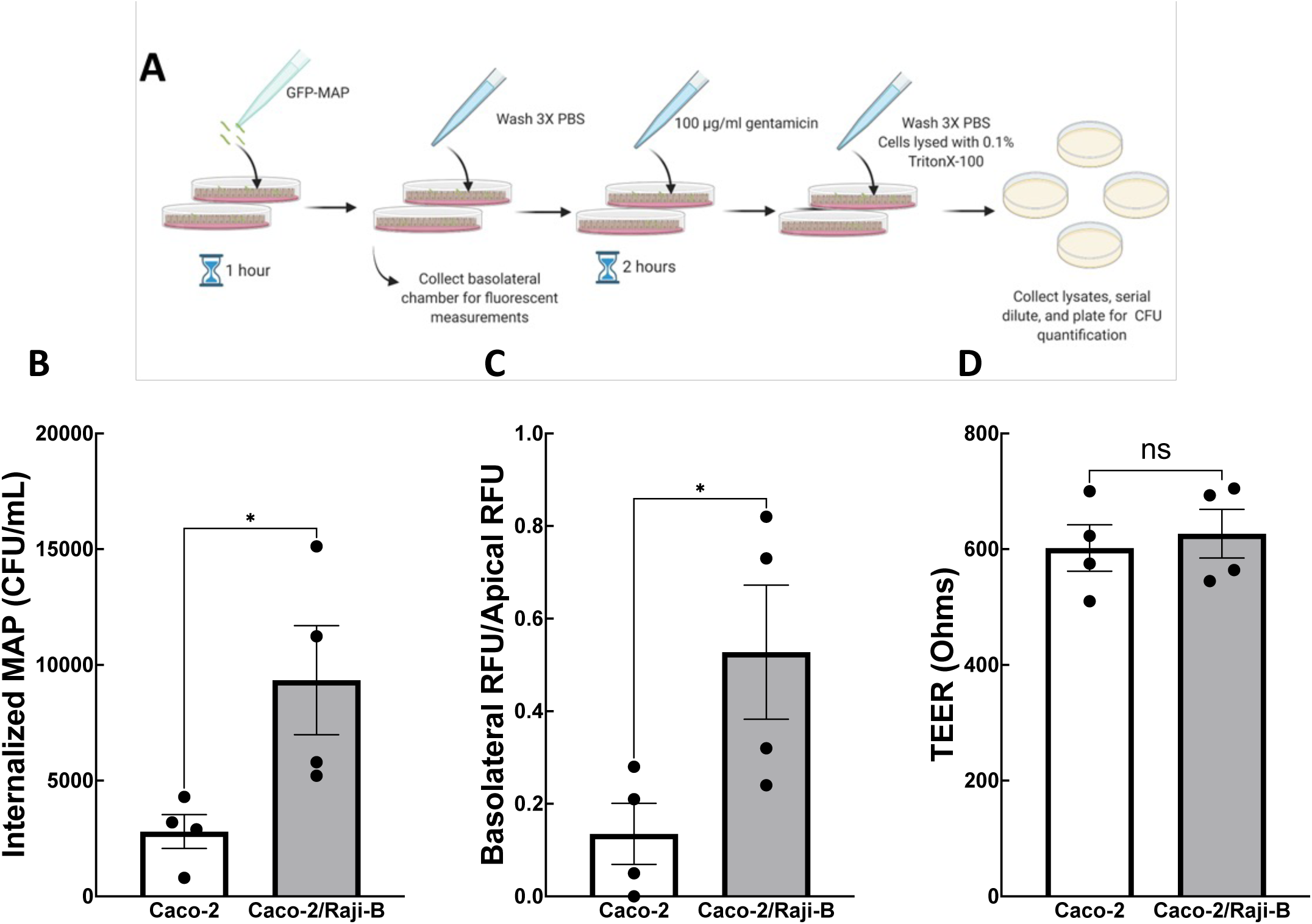
MAP internalizes and translocates through monolayers containing M cell-like cells to a significantly greater extent compared to mono-culture. Experimental schematic describing methods used to achieve results in B-E are displayed in (A). Internalized MAP was observed in both mono-culture and co-culture (B), however, the latter displayed a significantly higher quantity of internalized MAP. Basolateral MAP was quantified to assess MAP translocation across the monolayer (C). Results across 4 experiments show that the co-culture system had significantly more translocated GFP-MAP signal found in basolateral compartment compared to mono-culture system (C). To verify that this difference was not the results of differences in monolayer barrier function, TEER (D) was measured prior to GFP-MAP fluorescence measurements. Data are shown as means ± SEM. * P<0.05, n=4, with analysis performed via two-tailed Student’s t-test with Prism (GraphPad Software, La Jolla, CA).

### M cells within enteroid-derived monolayers exhibit apical expression of β1 integrin

We next sought to characterize the mechanism(s) responsible for MAP’s preferential targeting of M cells. Previous publications have implicated a fibronectin bridging system as the dominant mechanism of entry for other species of mycobacteria [40, 41]. Additionally, there is evidence to support the use of a fibronectin bridging system to allow for host uptake by MAP, however the host cell target of this system is unclear [42, 43]. Therefore, we wanted to evaluate any involvement of this system in MAP invasion of the host intestinal epithelium.

To do so, it was necessary to evaluate the presence of the key components of the fibronectin bridging system in enteroid-derived monolayers. We first assessed the expression of β1 integrin. We expected to see the presence of β1 integrin along the basolateral side of our monolayer, due to its involvement in attaching the basolateral membrane to extracellular matrix and adherence to the basement membrane below [44]. However, apical β1 integrin expression has been observed on M cells and shown to mediate binding and uptake [45]. We found that apical β1 integrin expression was absent in monolayers generated without RANKL treatment (Figure 10A). In contrast, β1 integrin expression (yellow) was prominent on the apical surface of RANKL-treated enteroid-derived monolayers, a signal that was co-localized with GP-2 (red), the mature M cell marker (Figure 10B-F). The apical expression of β1 integrin on GP-2 positive cells suggests that our RANKL-treated enteroid-derived monolayers contain M cells that express a key component required for bacteria to utilize a fibronectin bridge to facilitate binding.

**Figure 10:**
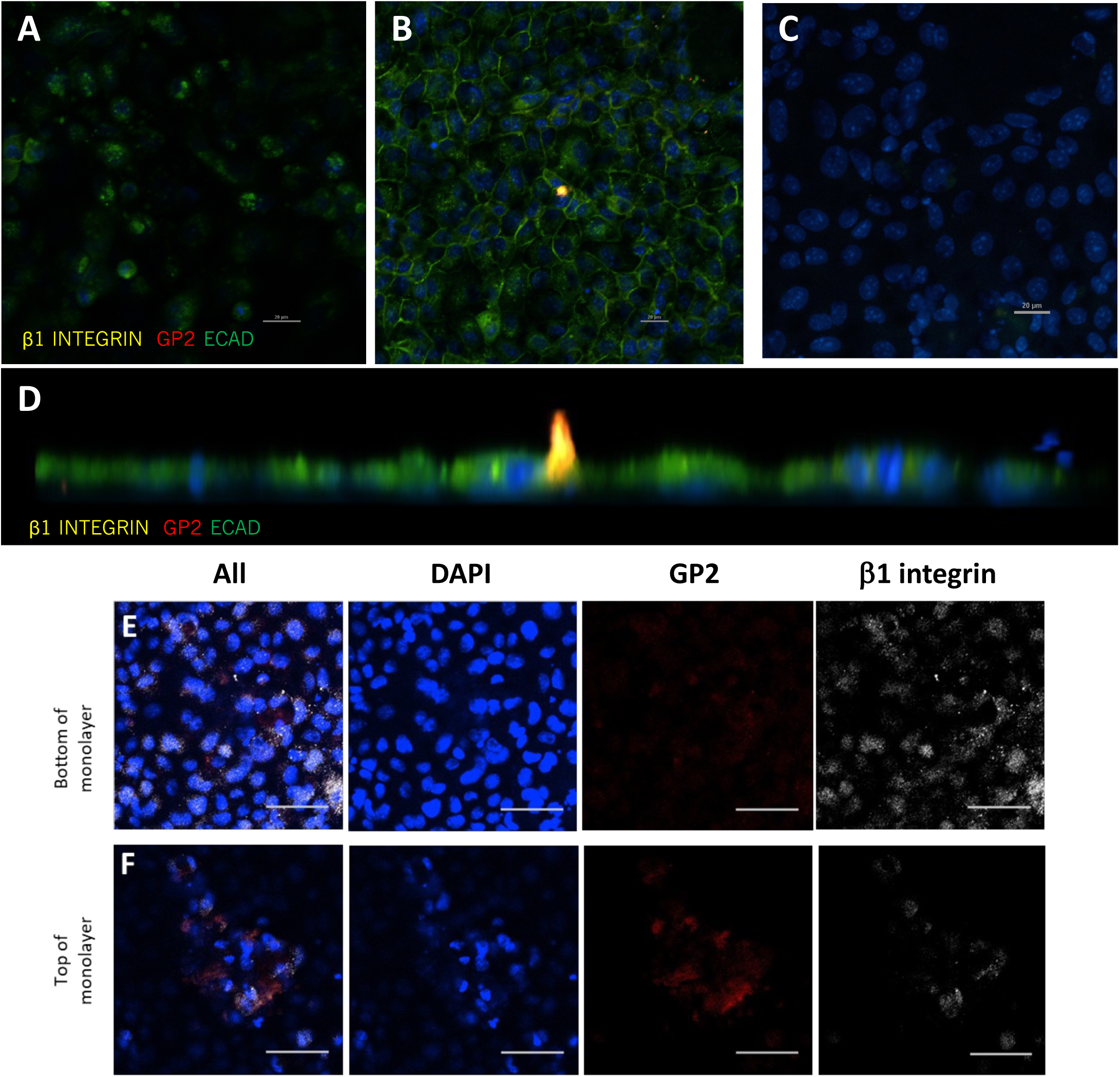
Mouse enteroid-derived monolayers exhibit β1 integrin expression on the apical surface that is co-localized with GP-2 expression in RANKL-treated monolayers. Mouse enteroid-derived monolayers with media alone (A) do not exhibit β1 integrin expression on their apical surface, however the addition of RANKL and the induction of M cell differentiation, as marked by GP-2 positive staining (B), results in expression of β1 integrin (yellow) co-localized with GP-2 positive staining (red), as evident in the z-stack 3D rendering (D). Specificity of antibody binding was evaluated via secondary alone antibody (C). Representative image from 3 separate experiments; DAPI (blue), ECAD (green), β1 integrin (yellow), GP-2 (red). β1 integrin expression is found on both the apical (E) and basolateral surface (F) of epithelial cells. To demonstrate this, representative images from a monolayer z-stack are shown here, with the top of the monolayer (E) and bottom of the monolayer (F) displayed to visualize this. Representative images of 4 separate experiments. Scale bar represents 50 µm. Images and z-stacks were captured on a Nikon-A1R confocal microscope.

### Fibronectin opsonization increases the number of MAP attached to and within M cells

With β1 integrin expression established in our enteroid-derived monolayer system, we next assessed whether the addition of fibronectin, the other key component of the fibronectin bridging system, would enhance MAP’s ability to adhere and invade M cells. In keeping with previous publications, MAP was incubated with PBS or fibronectin for 1 hour prior to exposure to M cell-containing enteroid-derived monolayers [43]. Fibronectin-opsonized MAP or MAP alone were incubated with monolayers for 30 minutes and then cells collected and stained for confocal immunofluorescence imaging. In keeping with our previous observations, GFP-MAP (green) was found within M cells (GP-2 positive staining regions; red) (Figure 11A, C).

**Figure 11:**
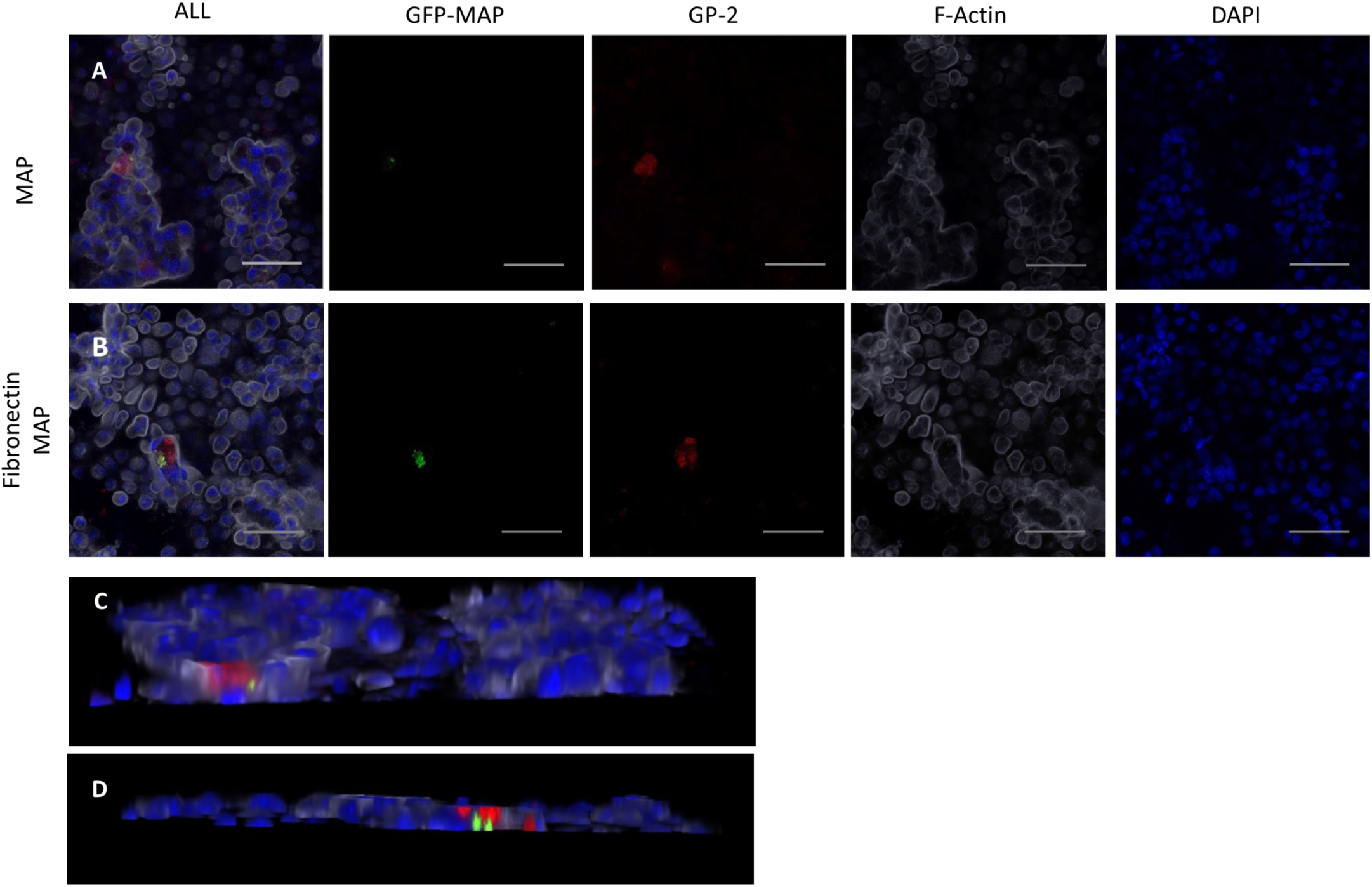
Fibronectin opsonization increases MAP uptake in M cells. RANKL-treated enteroid-derived monolayers were exposed to GFP-MAP (A) or GFP-MAP pre-incubated with fibronectin (FN; B) for 30 minutes prior to collecting monolayers for immunofluorescence imaing. As observed previously, GFP-MAP co-localized with GP-2 positive staining regions (A) as shown in the confocal image and z-stack rendering (C). However, FN-opsonized MAP resulted in higher numbers of discrete GFP-MAP, as seen in a representative confocal image (B) and z-stack rendering (D). Representative images from 5 separate experiments, scale bar represents 50 µm. Images and z-stacks were captured on a Nikon-A1R confocal microscope.

Incubating monolayers with fibronectin-opsonized MAP resulted in a greater signal intensity of GFP-MAP (green) and higher numbers of discrete GFP-MAP within M cells, as seen in a representative confocal image (Figure 11B) and z-stack rendering (Figure 11D). Upon enumeration of adherent and internalized MAP, we found that fibronectin-opsonization had no effect on enterocyte uptake of MAP, either in the number of MAP-positive enterocytes (Figure 12A) or in number of MAP found per enterocyte (Figure 12B). Our assessment of M cells revealed that fibronectin opsonization had no effect on the number of MAP-positive cells (Figure 12A), but increased the total number of MAP within individual M cells (Figure 12B) and the percentage of MAP adherent to M cells (Figure 12C). These data support the involvement of a fibronectin bridging mechanism to enhance MAP’s adherence and entry into M cells.

**Figure 12:**
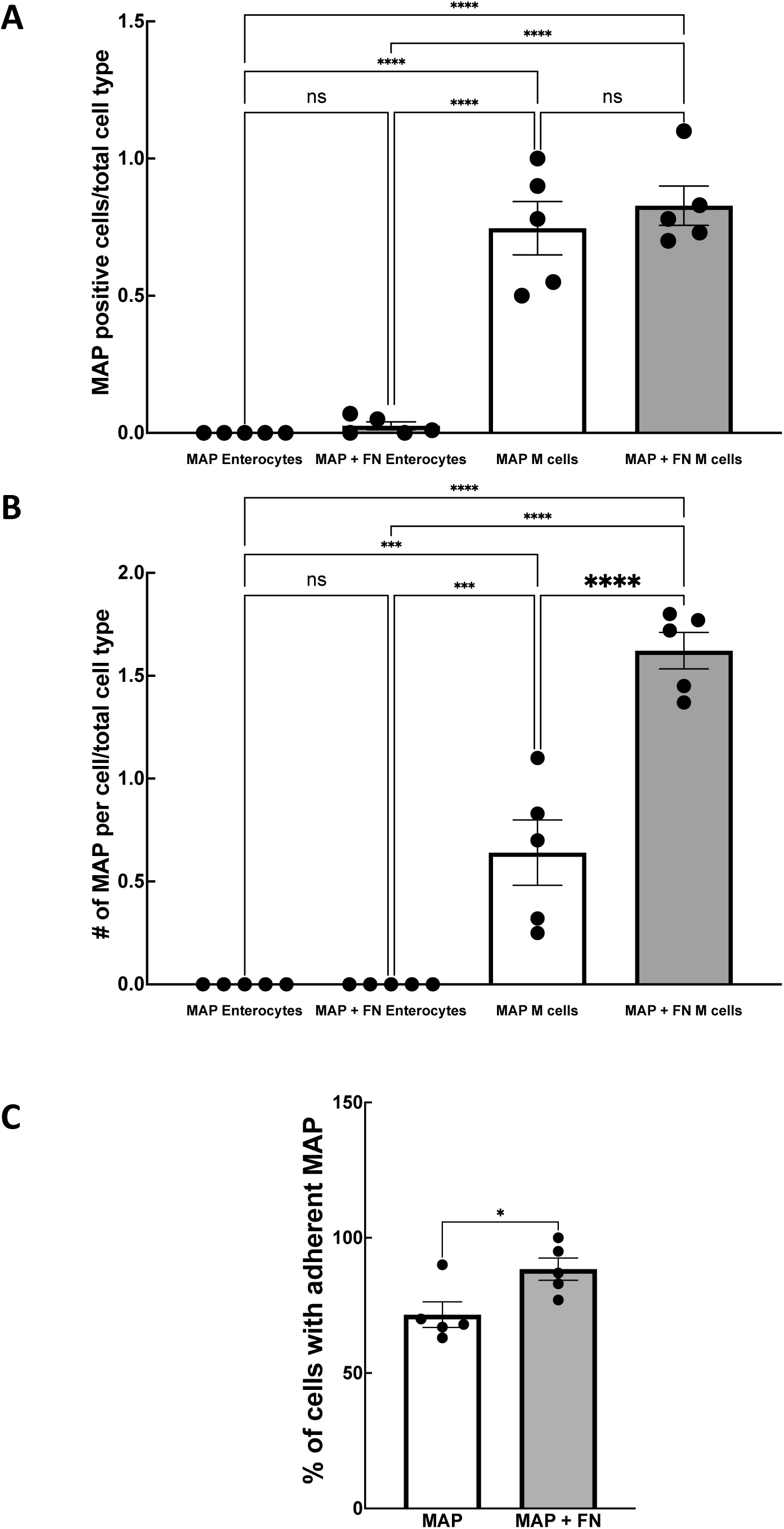
Fibronectin-opsonization of MAP results in no change in enterocyte uptake, but increases MAP uptake numbers observed per M cell. Quantitative analysis was performed from 5 unique experiments sets. MAP entry was observed in enterocytes, with no significant changes identified between fibronectin (FN)-opsonized and non-opsonized MAP (A-B). Number of MAP positive M cells (A) were analyzed for MAP and FN-MAP, no significant difference was observed between the two groups. MAP or FN-MAP number co-localizing with GP-2 positive stain was evaluated (B) and FN-MAP numbers in M cells were significantly increased compared to MAP alone. The percentage M cell adherent MAP was significantly higher in FN-treated MAP compared to MAP alone (C). Statistical tests used were two-tailed Student’s t-test, n=5 (** P<0.01) with values representing means ± SEM and normal distribution test performed via the Shapiro-Wilk test.

### Blocking the β1 integrin binding domain attenuated fibronectin-opsonized MAP invasion of M cells

Since we found that fibronectin opsonization increased MAP invasion of M cells, we next sought to assess the involvement of fibronectin-integrin interactions in the increased uptake observed. To test this, we pre-treated enteroid-derived monolayers with an RGD integrin-blocking peptide or a missense control peptide (GRAD) prior to exposure to fibronectin-opsonized MAP. Pretreatment with the RGD peptide significantly reduced the number of MAP positive M cells, when compared to monolayers pre-treated with the GRAD control peptide (Figure 13C-D). Interestingly, the RGD peptide had no impact on MAP within enterocytes (Figure 13E-F). Upon quantitative analysis of our images, we found that the RGD blocking peptide significantly reduced MAP within M cells, abolishing the M cell-tropism observed in our previous experiments (Figure 14A-B). Finally, we found that MAP adherence to M cells was significantly reduced by RGD-peptide treatment (Figure 14C) compared to monolayers treated with the GRAD control peptide. Taken together, these data suggest that fibronectin-β1 integrin interactions are key for the bacterium’s tropism for M cells and this process may constitute a potential target to reduce MAP infection in the GI tract.

**Figure 13:**
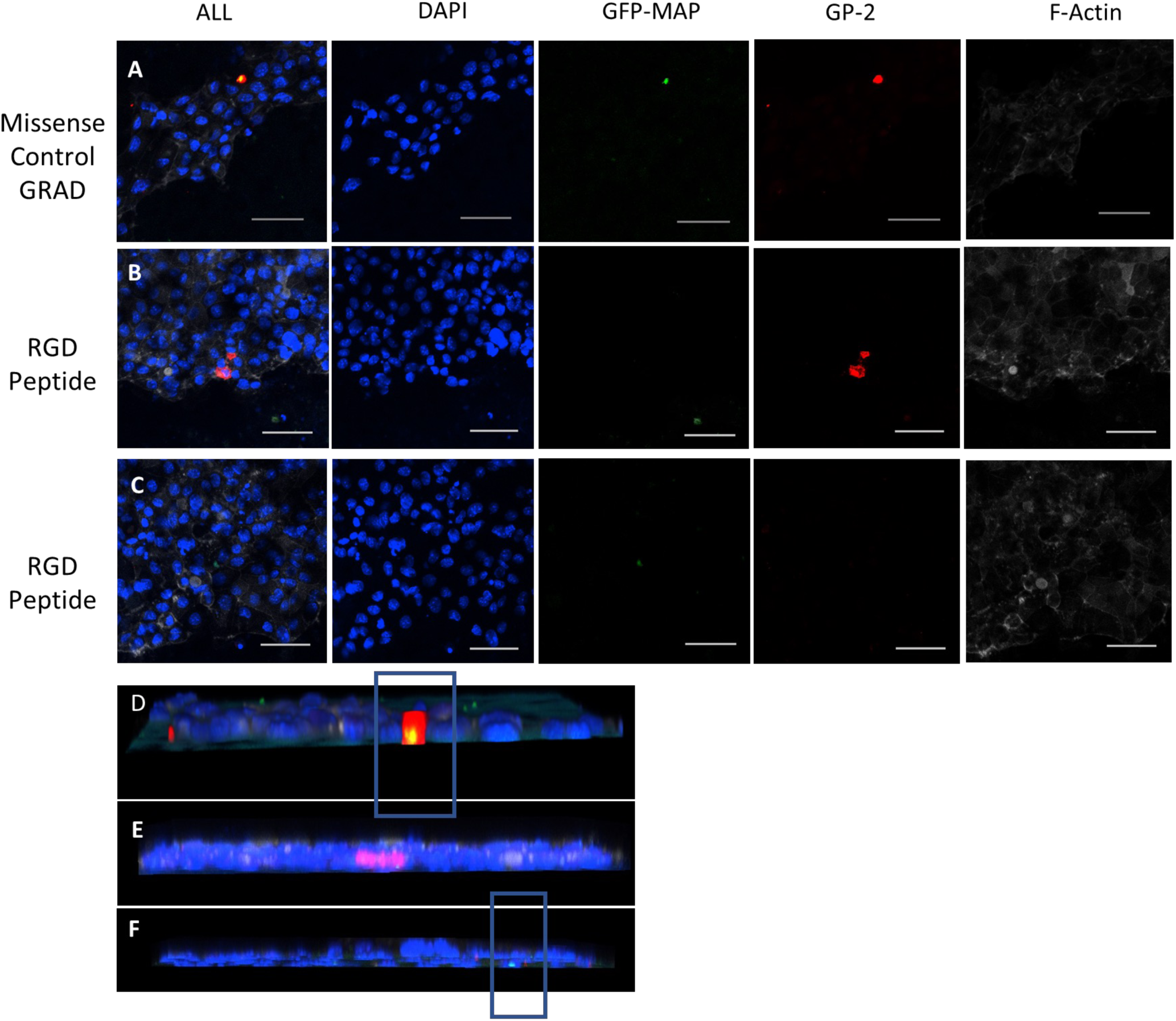
RGD peptide treated monolayers displayed decreased MAP uptake into M cells compared to missense peptide (GRAD). GFP-MAP positive signal within GP-2 positive signal was observed in missense control peptide GRAD-treated monolayers (A) with co-localization observed in z-stack 3D rendering (B). Pre-treating monolayers with the RGD-peptide significantly decreased M cell-associated MAP (C) as demonstrated by GP-2 positive signals in two separate DAPI-positive areas, both devoid of MAP. This is also visualized in the z-stack 3D rendering (D). RGD peptide-treated monolayers exhibited MAP uptake in regions devoid of GP-2 positive signal (enterocytes; E) and observed in z-stack 3D rendering (F). Representative images of 5 separate experiments. Scale bar represents 50 µm. Images and z-stacks were captured on a Nikon-A1R confocal microscope.

**Figure 14:**
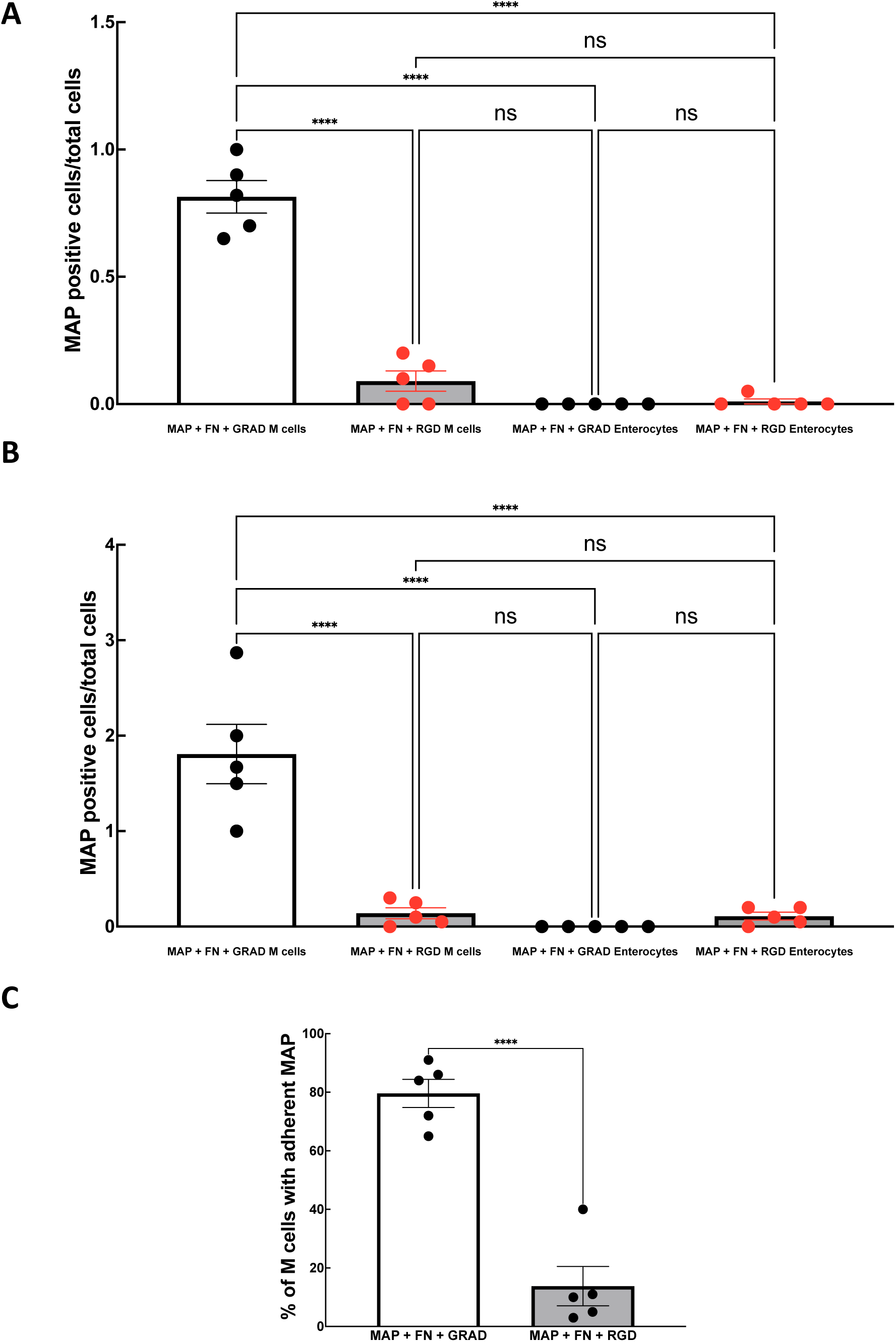
RGD peptide decreases MAP observed in M cells. Quantitative analysis was performed on 5 separate experiments. The number of MAP positive cells were enumerated with missense peptide (black data points) and RGD peptide-treated (red data points) (A, B). The number of MAP positive M cells was significantly reduced when monolayers were pre-treated with RGD peptide, compared to the control peptide (GRAD; A). MAP uptake in M cells was attenuated in RGD-treated group (MAP count per cell type; B), while MAP uptake in enterocytes is not affected. To evaluate percent of MAP adherent to M cells in fibronectin (FN) and RGD-treated monolayers, MAP adherence percentage was calculated for both group (C). Values depict means +/- SEM of 5 experiments (****P ≤ 0.0001). Mean values ± SEM were compared by one-way ANOVA with Tukey’s post-hoc test (GraphPad Software, La Jolla, CA) or Student’s t-test (C) with a P value of less than 0.05 considered statistically significant. To test for normal distribution, the Shapiro-Wilk test was used.

## Discussion

In the current study, we employed emerging enteroid-based approaches to test the hypothesis that M cells are the target for MAP attachment and invasion into the intestinal epithelium. Using mouse ileal enteroids seeded as monolayers, we demonstrated that MAP preferentially targets M cells, and this process involves interactions with apically expressed β1 integrins. Furthermore, MAP’s selective targeting of M cells could be enhanced by fibronectin-bridging, an effect that was attenuated by blocking β1 integrin-fibronectin interactions.

Most of the foundational studies seeking to characterize the interplay between MAP and the intestinal epithelium have used transformed cell lines, including non-intestinal cell models. One major limitation to these approaches is that these cell lines do not adequately model the cellular heterogeneity found within the intestinal epithelium because they do not differentiate into the various cell lineages. Furthermore, transformed cell lines exhibit significant alterations in their cellular physiology, including changes in metabolism, morphology, surface receptor expression, and thus do not reflect the attributes of the *in vivo* scenario that is being modeled [34, 42, 46]. In our studies, we generated enteroid-derived monolayers containing functional M cells to characterize MAP tropism and delineate the mechanism(s) associated with this selectivity. In keeping with the *in vivo* initial stages of MAP infection, a low concentration of MAP was added to the ileal enteroid-derived monolayer [47–49]. Previous studies using transformed cell lines have used a series of time-points to assess MAP uptake into epithelium. However, *in vivo* reports demonstrate that MAP uptake can occur within 30 minutes using a ruminant ileal loop model [26]. We observed rapid uptake of MAP within the intestinal epithelial monolayer, evident at the 30-minute timepoint. This uptake was seen in both M cells and enterocytes. Using confocal microscopy and a previously published protocol, we were able to enumerate the number of MAP-containing M cells and enterocytes in our experiments [50]. Surprisingly, while the number of enterocytes far outnumbered M cells within our monolayers, we repeatedly observed a greater frequency of MAP within M cells than enterocytes, even when normalizing for the proportion of the aforementioned cell types. Furthermore, complementary studies using the Caco-2/Raji-B co-culture approach demonstrated a similar degree of MAP M cell tropism in the context of a monolayer containing significantly more enterocytes compared to M cells. Collectively, these data suggest that MAP preferentially targets M cells. Furthermore, our study highlights the utility and value of enteroid-based approached for studying host-pathogen interactions, as these systems can provide the appropriate cellular heterogeneity (both diversity and quantity of cells) that best models the intestinal epithelium, allowing for researchers to identify and characterize mechanism(s) for attachment and invasion of enteric pathogens.

More recently, researchers have begun to develop enteroid approaches using bovine tissues allowing for the cultivation and long term maintenance of cultures from bovine small intestinal crypts [51]. While bovine enteroids retain morphology and the major functions of their digestive segment of origin [52], the field lacks the understanding of bovine M cell biology, including the pathways to drive their differentiation in culture, limiting the utility of this system to study MAP tropism at this time. Beyond this, there are limitations in the availability of reagents for use in bovine-based *in vitro* studies, including antibodies with specificity to bovine intestinal cell markers [51]. Those limitations aside, one recent publication investigated MAP uptake in bovine apical-out ileal enteroids and enteroid-derived monolayers [53]. While this group could detect genes associated with the presence of goblet cells, Paneth cells, enteroendocrine cells, and stem cells, they were unable to detect any expression profile suggestive of the presence of M cells. At this time, it is still not known if bovine M cells exhibit the same phenotypic markers used to identify this cell type in human or mice. However, reports have suggested that cytokeratin-18 or cyclophilin A might serve as markers of bovine M cells, but this has not been assessed in bovine enteroid systems [54, 55]. While bovine enteroids are likely the most appropriate model to study host-MAP interactions in the context of JD, it is apparent that more research is needed to identify bona fide M cell markers, and to demonstrate the existence of functional M cells within bovine enteroid-derived monolayers.

In contrast to bovine enteroids, previous studies have demonstrated functional M cells within human and mouse enteroids [30, 31]. Using the approaches reported by de Lau et al., we were able to generate 3D mouse ileal enteroids containing M cells that could be reseeded to form polarized monolayers, allowing us access to the apical aspect of the epithelium to subsequently incubate with MAP [30]. Previous studies have demonstrated that the GFP-MAP strain (K-10, pWES4) used our study can invade and infect mice both *in vitro* and *in vivo* [28, 48, 56–58].

Beyond this, the MAP K-10 strain has also been shown to cause infection and trigger paratuberculosis-like clinical signs in mice [48]. MAP has also been found in wood mice and it is thought that the lack of signs of overt clinical disease due to MAP infection is likely due to their short lifespan [59]. Together, these support the use of our well-characterized mouse enteroid-based systems to study MAP cell tropism, attachment and invasion.

In addition to identifying M cells as the primary target for MAP, we characterized key aspects of the mechanism responsible for this selectivity. Previous publications have implicated the fibronectin bridging system in the entry of MAP into enterocytes and M cells [43, 60, 61]. However, it is not known whether this system contributes to MAP M cell tropism. MAP has been shown to have express a fibronectin-attachment protein (FAP), which many species of mycobacteria utilize to attach to fibronectin *in vivo* [43, 62]. The MAP K-10 strain used in our studies has also been shown to bind to fibronectin [63]. *In vivo*, β1 integrins are expressed on the apical surface of M cells, and not found on the apical surface of enterocytes [64, 65]. After confirming the presence of apical β1 integrin expression within M cells, we sought to model the fibronectin bridging process by pre-treating MAP with fibronectin to effectively opsonize the bacteria prior to incubation with our enteroid-derived monolayers. While fibronectin opsonization had no effect on MAP attachment and entry into enterocytes, it significantly increased its adherence to and entry into M cells, suggesting a fibronectin bridging process could be at play. We next evaluated the role for β1 integrin in the mechanism underlying MAP uptake within M cells by pre-incubating our monolayers with an RGD peptide that acts to block the recognition sequence for integrin-fibronectin binding [61]. While the control missense peptide (GRAD) had no effect on MAP uptake in M cells, pre-incubation with the RGD blocking peptide nearly abolished MAP uptake into M cells, suggesting that fibronectin bridging is responsible for MAP’s M cell tropism. Interestingly, MAP uptake into enterocytes was not altered by the RGD peptide, suggesting that this process involves an alternate mechanism of adherence and eventual cell entry.

Moving forward, it will be imperative to characterize the exact mechanism that MAP utilizes to bind fibronectin. As discussed previously, the MAP K-10 strain used in our studies has also been shown to bind to fibronectin [63]. However, the discrete mechanism(s) responsible for this process remain to be elucidated. Several species of mycobacteria contain FAPs, including MAP [43]. FAP expression has been found to mediate attachment of both *M. avium* subsp. *avium* and *M. bovis* to host epithelium, suggesting a role for various FAPs in the attachment of mycobacteria to host tissues [45, 66]. Indeed, when FAP expression was attenuated in MAP strain 5781, equivalent levels of MAP were found in UAE-1 positive cells and enterocytes.

However a major caveat of this study is that UAE-1 is not a specific M cell marker, as it is also expressed on goblet cells [61]. Overall, while our data support that a fibronectin bridging mechanism is responsible for MAP’s selective targeting of M cells in our studies, understanding how the microbe interacts with fibronectin within the gastrointestinal tract to facilitate this process requires consideration in future studies.

## Methods

### Animals

All experiments involving mice were approved by the University of Calgary Animal Care Committee and all procedures followed their guidelines (protocol #AC20-0067). Male C57Bl/6 mice (12–25 weeks of age) were obtained from a colony from the Health Sciences Animal Resources Centre at the University of Calgary. Mice were housed in constant photoperiod (12 hours in light, 12 hours in dark) and temperature, with unrestricted access to laboratory chow and water.

### Enteroid generation and culture

Ileal crypts were isolated as described in previous publications [29, 67]. In brief, the latter third of the mouse small intestine was collected and placed in a tube with 30 mL PBS. Intestine segments were cleaned with PBS and cut into small pieces using mincing scissors. Tubes were then shaken and spun at 300 g for 30 seconds, resuspended in PBS and spun two additional times, also at 300g for 30 seconds. Tissue was then resuspended in 30 mL of crypt isolation buffer (2 mM EDTA in PBS + 43.4 mM sucrose + 0.5 mM DTT) and incubated at 4°C with gentle circular shaking (speed 12-20 rpm). After 30 minutes, the tissue was spun at 300 g for 30 seconds and the supernatant was removed. At this point, 10 mL cold PBS was added to tissue, the mixture resuspended 6 times and then the supernatant was placed into a fresh 50 mL tube.

This process was repeated 5 times, and each time the supernatant was placed into a new 50 mL tube. Supernatants were strained through 100 μm strainer, and the flow-through collected in a new tube, and centrifuged at 300 g for 5 minutes at 4°C. Supernatant was then removed via aspiration. Pellet was then resuspended in Matrigel (354230, Corning). Media was generated as described previously [29]. Enteroids were passaged weekly or once cell debris accumulated within the inner lumen (as outlined in [29, 67]). All experiments were performed between passages 3-30.

### Composition of mouse enteroid media

Conditioned media from L-WRN cells (CRL-3276; ATCC) was generated as previously described, where media contained Wnt-3A, R-spondin 3, and Noggin [68]. Conditioned media was then diluted in basal crypt media containing Advanced DMEM/F12 (ADF) (12634028; Gibco) with Glutamax (2 mM; GIBCO, ThermoFisher, Waltham, MA), N-acetyl cysteine (NAC; 1 mM; Sigma-Aldrich), HEPES (10 mM), penicillin (100 units/ml), streptomycin (0.1 mg/ml), N2 Supplement (1×, Invitrogen), and B27 supplement (1×, Invitrogen). Final media was generated by combining L-WRN media and basal crypt media at a 1:1 ratio. After the first feeding post passaging, Y-27632 (ROCK inhibitor, 10 µM) was added. For each 3D enteroid media change, primocin (0.1 mg/ml) was added to prevent contamination. For monolayers, NAC was omitted to promote enteroid monolayer growth and human epidermal growth factor (hEGF; 50 ng/ml; Austral Biologicals, San Ramon, CA) was added. Additionally, each monolayer feeding contained both Y-27632 (10 µM), and primocin (0.1 mg/ml) to supplement final complete media composition.

### M cell differentiation in enteroid cultures

Ileal enteroids were generated as described previously [29, 67, 69]. To induce the differentiation of M cells, 100ng/mL recombinant mouse RANKL (315-11; Peprotech) was added on day 3 of enteroid culture; control plates were maintained in media without RANKL, following previous publications [30, 70]. On day 6, enteroids were harvested, and collected for immunofluorescent staining. RNA was also collected for qRT-PCR assessment of M cell markers.

To generate 2D enteroid-derived monolayers containing M cells, 3D ileal enteroids were treated with 100ng/mL RANKL for 3 days, as described above. On day 6, enteroids were dissociated and plated as monolayers on 3µm transparent membrane (PET) (662630; Greiner bio-one) pre-coated with Matrigel [29]. Monolayer media was supplemented with 100ng/mL RANKL in the basolateral chamber for experiments requiring differentiated M cells. Media was changed daily, and monolayer confluence was assessed by measuring transepithelial electrical resistance (TEER). Differentiated M cells were detected through confocal microscopy (Nikon-A1R) and immunofluorescence staining for glycoprotein-2 (GP-2).

### Caco-2 and Raji-B cell culture

The human colorectal adenocarcinoma cell line Caco-2 (HTB-37; ATCC) was grown in DMEM with 20% FBS (F1051-500mL; Sigma), 1% sodium pyruvate (S8636-100mL; Sigma-Aldrich), penicillin–streptomycin (100ug/mL, P4333; Sigma) 1% non-essential amino acids (11140-050; Thermofisher), L-glutamine (G7513; Sigma) and 1mM HEPES (15630-080; Gibco). Human Burkitt lymphoma cell line, Raji-B (CCL-86; ATCC) was grown in RPMI with 10% FBS and L-glutamine (G7513; Sigma). All cells were maintained in a humidified 37°C incubator and 5% CO_2_ for the duration of experiments.

### Caco-2/Raji-B Co-Culture

As a complementary experimental system to study M cell biology, Caco-2/Raji-B co-cultures were performed as described previously [35, 71]. In brief, Caco-2 cells (1×10^6^ cells/mL) were seeded on Matrigel-coated transwells and TEER assessed daily. Media was changed every other day until day 14, at which point 5×10^5^ cells/mL Raji-B cells were added to the basolateral compartment of the Transwell. Control monolayers (without differentiated M cells) were cultured in the absence of Raji-B cells. Media was changed daily for three additional days or until TEER was reduced by 10% (in Caco-2/Raji-B co-culture monolayers). At this point, experiments were performed using Caco-2/Raji-B or control monolayers. Differentiated M cells were detected through confocal microscopy and immunofluorescence staining for sialyl Lewis A antigen (SLAA; 2.5µg/mL, 2194142; Novus Biologicals)).

### MAP Culture

Mycobacterium avium subspecies paratuberculosis MAP-K10 (ATCC BAA-968; referred to as GFP-MAP) expressing green fluorescent protein was used for all experiments. This strain was originally developed by Parker and Bermudez, by transforming the wild-type K10 MAP strain with a recombinant plasmid (pWES-4) expressing GFP and a kanamycin resistance gene; insertion of the plasmid has not been shown to alter the virulence of MAP [72]. For GFP-MAP culture, Middlebrook 7H9 medium was used (Difco Laboratories, USA) and supplemented with 0.2 % glycerol, 10% oleic acid-dextrose-catalase (OADC) and mycobactin J (2mg/L) (Allied Monitor) and grown at 37°C in a shaking incubator at 120 RPM. Bacteria were used when the OD_600_ was 0.6-0.7. Cultured GFP-MAP concentration (bacteria/mL) was determined by pellet weight following the methods published by Hines and others [73]. In brief, GFP-MAP was centrifuged at 3000 x g for 10 minutes. The pellet was then cleaned with PBS twice and allowed to dry. The pellet was then weighed, using the conversion where 1 mg is equivalent to 1 × 10^7^ CFU. CFU counts were then confirmed via serial dilution on 7H11 plates, where 7H11 plates were supplemented with 2 mg/L mycobactin J (Allied Monitor), 10% OADC, 10µg/mL vancomycin (V2002, Sigma) 20µg/mL amphotericin B(A2942-20ML, Sigma-Aldrich), and 30µg/mL nalidixic acid (N8878, Sigma) to prevent culture contamination [74, 75]. GFP-MAP cultures used in all experiments were plated on blood agar plates to ensure the absence of other bacteria.

### RNA extraction and Quantitative real-time PCR

Matrigel-embedded enteroids were washed three times with PBS, and 200 µL Trizol (T9424-200 mL; Sigma Aldrich) was added to each Matrigel dome in a 24 well plate. The contents of 4 separate replicate domes were pooled for each experimental condition for 3D enteroids and 4 replicate Transwells were pooled for each experimental condition in mouse enteroid monolayers. Chloroform (200 µL) was added for every 1 mL Trizol (T9424-200 mL; Sigma) and the solution was shaken vigorously by hand for 15 seconds. After incubation at room temperature for 5 minutes, samples were centrifuged at 12,000 rpm for 15 minutes at 4°C and the aqueous phase was removed. RNA was isolated using the RNeasy minikit (74106; Qiagen) according to the manufacturer’s instructions. cDNA was generated using the QuantiTect Rev. Transcription Kit (205314, Qiagen) according to the manufacturer’s instructions. Real-time PCR kit, PerfeCTa® SYBR® Green FastMix®, ROX™ (101414-284; VWR) was used for all PCR assays on the StepOnePlus™ Real-Time PCR System. All reactions were run in duplicate or triplicate.

Validated primers from Qiagen were used to quantify the expression of target genes in control and RANKL-treated 3-D/2-D enteroid cultures (Table 1.0). Relative changes in mRNA expression were determined by comparative threshold (C_t_) analysis using ΔΔCt method of relative quantification and are presented as the mean ± SEM of independent experiments.

**Table 1.0:** Primers used for qPCR; validated RT primers obtained through Qiagen. In all samples, β-actin (Actb) was used as the endogenous control.

| *Gene* | *ABRV* | *Refseq ID* | *Company* | *Catalogue No.* |
| --- | --- | --- | --- | --- |
| *Spi-B transcription factor* | *Spi-B* | NM_019866 | Qiagen | PPM04709A |
| *Glycoprotein 2* | *GP-2* | NM_025989 | Qiagen | PPM29342A |
| *β-actin* | *Actb* | NM_007393 | Qiagen | PPM0245B |

### Bead Uptake Assay

To assess the functional properties of differentiated M cells in enteroid-derived monolayers or within Caco-2/Raji-B co-cultures, active uptake of fluorescent polystyrene beads (1×10^5^ 1 µm; F13083; Invitrogen) was assessed. Beads were added to the apical side of the monolayers at various days post-plating (after 18 days of growth for Caco-2/Raji-B co-cultures) and incubated for 1 hour, according to a previously published protocol [31]. Monolayers were then washed three times in PBS to remove any remaining microparticles and immunofluorescent staining was performed. Confocal microscopy was performed to evaluate bead and cell marker co-localization.

### Analysis of Bead and GFP-MAP Uptake

To quantify bead uptake, the immunofluorescent z-stacks were analyzed using a previously published approach [31], based on FIJI [76]. The number of enterocytes per image was quantified by enumerating E-cadherin positive cells, whereas M cells were quantified by counting GP-2 positive cells. To visualize and quantify beads the threshold of the red channel (emission from beads) was automatically set using the Otsu method, as described previously [31], until all areas positive for bead-associated signal were clearly defined.

Attached and internalized beads associated with enterocytes (E-cadherin positive) and M cells (GP-2 positive) were quantified in z-stacks. This approach enumerated (per field of view) total cells, total M cells, total enterocytes, total bead counts, total beads attached or internalized into M cells or enterocytes and the total number of bead-positive M cells and enterocytes. These data allowed us to calculate the percentage of bead-positive (adherent or internalized) M cells and enterocytes for statistical purposes.

### MAP exposure

Enteroid-derived monolayers were exposed to GFP-MAP through its addition to the apical compartment of the transwell. GFP-MAP was added at 5×10^5^ CFU/mL, a concentration that represents low level of MAP exposure *in vivo* [47, 48, 77]. GFP-MAP-exposed monolayers were then placed in a 37°C incubator for 30 minutes, then washed three times with ice-cold PBS prior to processing for immunofluorescence imaging.

### Analysis of MAP Uptake

To quantify the immunofluorescent z-stacks, the data collected from MAP co-culture with mouse monolayers were analyzed based on a published report [50]. Images were captured on Nikon-A1R confocal microscope in the LCI Laboratory (RRID SCR_024748) at the University of Calgary. To avoid bias, fields-of-view were visualized using the GFP-MAP signal alone. Once MAP was visualized, a z-stack was captured using the full set of lasers to excite additional fluorescent labels within the samples. Controls for each experiment included a no MAP control, a monolayer without RANKL (without M cells), and a secondary antibody only control. These controls were measures of non-specific signaling.

Images were saved as ND2 (Nikon) files and imported into FIJI for downstream analysis [76]. The number of epithelial cells per image was calculated by measuring the number of nuclei detected from the DAPI channel. To quantify the number of GFP-MAP particles, the emission signal was automatically thresholded using the Otsu method, following a previous publication [50]. Similarly, the M cells were identified in the GP-2 channel by using Otsu thresholding.

Numbers of invading MAP were enumerated through visualizing the z-stacks using 3D rendering using Nikon-A1R Elements. Cells were considered positive for MAP adherence and/or internalization only if GFP-MAP fluorescence signal was observed within the outline of the cell or within the cell lining (within or overlapping with ECAD or GP-2).

This method allowed for quantification of total cells, M cell number, non-M cell number, MAP count, MAP attached or internalized in M cells, MAP per M cell, MAP attached or internalized in non-M cells, and adherent MAP. At this point, we could evaluate percent MAP^+^ cells, number of MAP invading per cell type, and adherent MAP.

### Fibronectin Opsonization Assay

For fibronectin experiments, bovine fibronectin (2µg/mL; FC014; Sigma-Aldrich) was added to MAP culture for one hour prior to experiments, following a previous publication [61]. GFP-MAP alone was also incubated at 37°C for 1 hour as a control. Bacterial suspensions were then added to 2D ileal monolayer system for 30 minutes.

### Integrin Blocking Peptide Assay

For integrin blocking experiments, RGD (GRGDSP; SCP0157-5MG; Millipore Sigma) and control (GRADSP; SCP0156-1MG; Sigma Aldrich) peptides were resuspended in ANT buffer (10 mM ammonium acetate, 0.85% sodium chloride, 0.05% Tween 20, pH 6) to a concentration of 5 mM, following a previous publication by Secott and others (2004). Peptides were added to monolayers 30 minutes prior to MAP exposure and incubated for 30 minutes to allow for MAP exposure on the mouse ileal enteroid-derived monolayer system.

### Immunofluorescence

After each exposure protocol was completed, 3D enteroids and 2D ileal enteroid-derived monolayers were washed twice with PBS prior to 4% PFA treatment for 15 minutes, as described in a previous publication [29]. The wells were washed with PBS three additional times and blocked with 5% NGS for an hour at room temperature or overnight at 4°C. Wells were then washed three times with 0.1% PBS-T and cells were permeabilized with 1% Triton-X (#9002-93-1; BioPharm). Primary antibodies were added and incubated at room temperature for an hour (list of primary antibodies can be found in Table 2.0). Cells were washed three times with 0.1% PBS-T and incubated with secondary antibody (1:1000) for an hour (list of secondary antibodies can be found in Table 3.0). Next, cells were washed three times with 0.1% PBS-T and DAPI (1:50,000; D1306; Invitrogen) was added for 30 minutes. Finally, cells were washed with PBS three times and mounted using Prolong Gold Antifade Mountant (P36970; Thermofisher).

**Table 2:** Primary antibody concentrations, catalogue number, and company.

| <u>Antibody</u> | <u>Concentration</u> | <u>Catalogue #</u> | <u>Company</u> |
| --- | --- | --- | --- |
| *Mouse GP-2* | 2.5µg/mL | D278-3 | MBL |
| *Human GP-2* | 1:200 | D277-A48 | MBL |
| *F-actin* | 1:200 | A22287 | Thermofisher |
| *E-Cadherin* | 1:200 | 610181 | BD Bioscience |
| *β1 integrin antibody* | 1:500 | ab179471 | Abcam |
| *SLAA* | 5µg/mL | 2194142 | Novus Biologicals |
| *ZO-1* | 1:100 | 33-9100 | Invitrogen |
| *RANK* | 1:100 | sc-390655 | Santa Cruz |

**Table 3:**
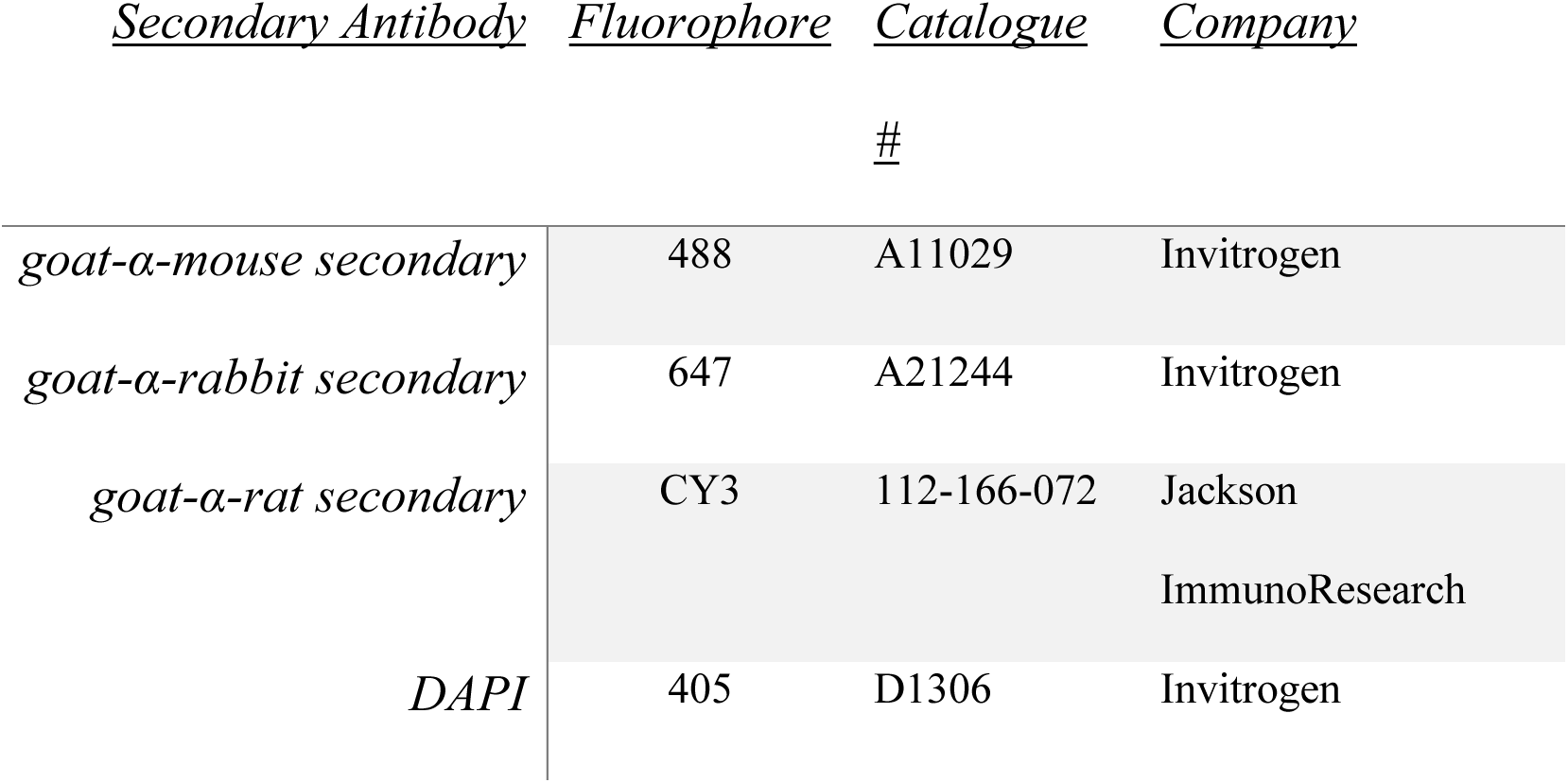
Fluorescent secondary antibodies and DAPI concentrations, catalogue number, and company.

For 3D culture, enteroids were mounted on cavity slides to prevent flattening of the 3D structure of the enteroids. For 2D culture, monolayers were stained following the methods described above, and imaged in the transwell directly after staining. For immunofluorescence and differential interference contrast (DIC) confocal imaging, a Nikon-A1R confocal microscope was used.

### Antibodies

3D enteroids cultured with or without 100ng/mL RANKL were stained for GP-2 (2.5µg/mL; D278-3; MBL), F-actin (1:200, Thermofisher, Cat #A22287), ECAD (1:200; 610181; BD Bioscience), RANK (1:100; sc-390655; Santa Cruz) and DAPI (1:50,000; D1306; Invitrogen). For 2D epithelial monolayer studies, GP-2, F-actin, and ECAD were also utilized. For Caco-2 monolayers, sialyl Lewis A Antibody (SLAA) (2.5µg/mL, 2194142; Novus Biologicals) was also used. Table 2.0 describes the list of primary fluorescent antibodies used in this investigation. Table 3.0 describes the secondary antibodies used in this study. All secondary antibodies used in this investigation were used at 1:1000 dilution.

### Fluorescence Intensity Readings

Fluorescent signal was measured in Greiner black polystyrene 96 well plates with the SpectraMax i3x plate reader (Molecular Devices, San Jose, CA) using 100 µL of supernatant from the apical or basolateral chamber of transwells in various experiments. For fluorosphere translocation assays, fluorescence signal was quantified; a standard curve was used to determine the concentration of translocated beads from the signal. By comparing basolateral chamber signal to the standard curve, we could calculate translocated beads/mL in mono- and co-culture systems. For GFP-MAP translocation assays, GFP RFU signal from the basolateral chamber was compared to GFP RFU signal from the GFP-expressing MAP that was added to the apical chamber at the start of the experiment.

### Trans-Epithelial Electrical Resistance (TEER) Measurements

2D ileal enteroid-derived monolayers and Caco-2/Raji-B co-cultures were grown in 500µL apical and 500µL basolateral conditioned media. Culture media was replenished 30 minutes prior to measuring TEER with a Millicell ERS-2 voltohmeter (Millipore), following the manufacturer’s instruction. Each insert was measured three times and averages values were corrected compared to the background TEER, measured using the Matrigel-coated Transwell alone (no cells were plated on these Transwells). This value was then displayed in Ohms (Ω) corrected to baseline Matrigel-coated Transwell membrane or raw Ω values, with Matrigel-coated Transwell Ω values indicated by a dashed line.

### Statistical Analysis

Statistical analysis was performed using GraphPad Prism (GraphPad Software, La Jolla, CA). Data are presented as means with bars representing standard errors of the mean (SEM). Prior to statistical analysis, all data were assessed for normal distribution (Shapiro-Wilk test). Data deemed to exhibit a normal distribution were analyzed using parametric approaches. (unpaired Student’s *t*-tests to compare two groups or one-way and two-way analysis, an analysis of variance (ANOVA) was used followed by a Tukey or Bonferroni’s *post-hoc* multiple comparisons test).

## Acknowledgements

We would like to acknowledge the support and guidance of Dr. Lucy Swift, of Senior Optical Imaging Specialist, the Live Cell Imaging Laboratory (RRID SCR_024748) funded by the Snyder Institute for Chronic Diseases.

